# Development of the novel amylin receptor activators with nanomolar potency by peptide mutagenesis

**DOI:** 10.1101/2024.07.31.606116

**Authors:** Sangmin Lee

**Affiliations:** Department of Medicinal Biotechnology, College of Health Science, Dong-A University, Busan, Republic of Korea, 49315

**Keywords:** Affinity and potency/amylin peptide hormone/mutagenesis/obesity

## Abstract

Amylin peptide hormone activates amylin receptors in brains and controls blood glucose and appetite. An amylin receptor activator pramlintide was developed for diabetes treatment. Currently, the amylin receptor activator with once-weekly injection has been tested for body weight reduction to treat obesity. Human amylin peptide was reported to form aggregates, while rat amylin has been shown soluble in aqueous solution. Here, multiple peptide activators for human amylin receptors were developed by introducing comprehensive mutagenesis to rat amylin peptide. The rat amylin peptide C-terminal fragment is known for interacting with human amylin receptor extracellular domain. The rat amylin peptide C-terminal fragment with eleven amino acids was used to screen for affinity-enhancing mutations. Up to twelve mutational combinations were found to significantly increase peptide affinity for amylin receptor extracellular domains by over 100-fold. Using these affinity-enhancing mutations, three representative rat amylin analogs with thirty-seven amino acids were made to test the potency increase for amylin receptor activation. All three mutated rat amylin analogs showed significant potency increases by 5- to 10-fold compared to endogenous rat amylin. These mutated peptide activators also showed higher potency for human amylin receptor activation than a clinical drug pramlintide. These amylin receptor activators developed in this study can be useful for the drug development targeting diabetes/obesity treatment.

## 1. Introduction

Obesity is a common and serious disease worldwide with limited pharmacological options. Obesity is a leading contributor to type II diabetes development [1] and it also induces multiple complications such as cardiovascular diseases and joint pain [2]. Peptide hormone analogs that activate glucagon-like peptide 1 (GLP-1) receptors were recently approved for obesity treatment. GLP-1 receptor activators originally have been approved for type II diabetes by increasing insulin secretion [3]. Among them, liraglutide and semaglutide were reported to reduce body weight [4,5] and most recently once-weekly injection of semaglutide was approved for obesity treatment [6].

In addition to GLP-1, two more peptide hormones have shown potential for obesity treatment: glucose-dependent insulinotropic polypeptide (GIP) and amylin. GIP stimulates insulin secretion and is believed to reduce appetite. Once-weekly injection of a GIP and GLP-1 receptor dual agonist tirzepatide showed promising results for obesity treatment [7] and its use for body weight control was recently approved in US.

Amylin is secreted from pancreas with insulin and amylin receptor activation controls blood glucose and body weight [8]. An amylin analog pramlintide has been approved for diabetes treatment [9]. Recent clinical trials with once-weekly injection of an amylin receptor activator cagrilintide [10] alone and cagrilintide combination with a GLP-1 receptor agonist semaglutide showed a significant 10 to 17% body weight reduction [11,12].

The amylin receptor is the heterodimer complex of the calcitonin receptor and an accessary protein called receptor activity-modifying protein (RAMP) [13,14]. There are three types of RAMP in humans (RAMP1–3), and the calcitonin complex with each of RAMPs constitutes amylin receptor 1–3. Recent cryo-electron microscopy (cryo-EM) structures of the amylin receptors clearly showed the binding mode of peptide hormone amylin and calcitonin for their receptors [15]. The N-terminal fragment of amylin interacts with the transmembrane domain of the calcitonin receptor. In contrast, the C-terminal fragment of amylin interacts with the extracellular domains of the calcitonin receptor and RAMP [16].

Human amylin peptide was reported to form aggregates that were visualized as amylin fibrils by cryo-EM [17]. However, rodent amylin has a different amino acid composition compared to human amylin and it has been shown soluble in aqueous solution. Especially, rat amylin has been commonly used for animal studies and for cell signaling assay to activate amylin receptors. This study focuses on the design of potent rat amylin analogs that may enhance anti-obesity efficacy through amylin receptor activation. The hypothesis for this study was that introducing mutations to the peptide fragments can produce novel peptide ligands with altered (preferentially enhanced) affinity and potency for the amylin receptors. The primary endpoints of this study were to find affinity-enhancing mutations by at least 10-fold and to develop potency-enhancing peptide activators by 3-fold when compared to endogenous rat amylin. This study measured the binding affinity of the rat amylin peptide C-terminal fragment for purified human amylin receptor extracellular domains with fluorescence polarization peptide-binding assay. Comprehensive mutagenesis was introduced to the rat amylin peptide C-terminal fragment to find affinity-enhancing mutations. The resulting affinity-enhancing mutations were used for potency enhancement for human amylin and calcitonin receptor activation.

## 2. Materials and Methods

### 2.1. Reagents

Dulbecco’s Modified Eagle Medium (DMEM) including 4.5g/L glucose, L-glutamine, and sodium pyruvate was purchased from Corning (Mediatech, Inc., Manassas, VA) to culture mammalian cells. The mixture of non-essential amino acids (NEAA, 100X) was purchased from Lonza (Basel, Switzerland). For mammalian cell culture, fetal bovine serum (Cat.# F2442) was purchased from Sigma-Aldrich (St. Louis, MO). NEBuilder® HiFi DNA Assembly Master Mix and several restriction enzymes were purchased from New England Biolabs (Ipswich, MA) and were used for cloning plasmid DNA constructs. 4–12% Mini-PROTEAN TGX Stain-Free Precast Gels were purchased from Bio-rad (Hercules, CA). All other reagents were purchased from Sigma-Aldrich, unless otherwise noted.

### 2.2. Cell lines and bacteria cells used

Human embryonic kidney (HEK) 293T cells and HEK293S GnTI^-^ cells were purchased from ATCC (Manassas, VA) to express calcitonin and amylin receptor extracellular domains. HEK293T cells were also used to assess calcitonin and amylin receptor activation by using bioluminescence resonance energy transfer (BRET) assay. This BRET assay was well-established with HEK293 cells in a previous study [18] and using one cell-line for BRET assay was assumed reasonable. The cell lines used for this study have been authenticated from the provider ATCC and the cells with the passage number less than 30 were used for this study. One Shot™ OmniMAX™ 2 T1R Chemically Competent *E. coli* cells (Cat.# C854003) were purchased from Invitrogen for DNA cloning.

### 2.3. Expression plasmids

Previously reported pHLsec-based vectors were used to express calcitonin and amylin receptor extracellular domains as secreted proteins [19]. The following plasmid DNA constructs used in this study were previously described [20,21]: pHLsec/hCTR.34–141-H_6_ (H-pSL003), pHLsec/hRAMP1.24–111-(GSA)_3_-hCTR.34–141-H_6_ (H-pSL005), pHLsec/hRAMP2.55–140-(GSA)_3_-hCTR.34–141-H_6_ (H-pSL006), and pHLsec/hRAMP3.25–112-(GSA)_3_-hCTR.34–141-H_6_ (H-pSL001). The following constructs for bioluminescence resonance energy transfer (BRET) assay that assesses G protein recruitment to G protein-coupled receptors [18] were generously gifted from Dr. Nevin Lambert (Medical College of Georgia): NES-Venus-mGs (H-pSL028) and Nluc-N1 (H-pSL033). The cDNA sequence of full-length human calcitonin receptor was purchased from CDNA.org (Cat.# CALCR00000, H-pSL053). The cDNA sequences of human receptor activity-modifying protein 1 (RAMP1) (NM_005855.4, H-pSL056), RAMP2 (NM_005854.3, H-pSL042) and RAMP3 (NM_005856.3, H-pSL043) were purchased from Genscript (Piscataway, NJ, USA). The DNA construct of the calcitonin receptor C-terminally fused with NanoLuc luciferase [22] (Nluc_FL_CTR (H-pSL054)) was generated for the current study using DNA Assembly reaction. Coding sequences of the above expression vectors were confirmed with Sanger sequencing performed by Psomagen (Rockville, MD). DNA expression plasmids were extracted and were purified from bacterial cells with NucleoBond® Extra Midi Plus kit (Macherey-Nagel, Germany) and they were stored at -20°C until their use for transient transfection into mammalian cells.

### 2.4. Expression and purification of RAMP ECD-CTR ECD fusion proteins

The general expression procedure of the calcitonin receptor (CTR) extracellular domain (ECD) alone and RAMP1, -2, or -3 ECD fused with CTR ECD as functional amylin receptor ECDs was previously reported [21,23]. Briefly, the amylin receptor ECDs were expressed from HEK293T or HEK293S GnTI^-^ cells with transient transfection. The expressed receptor proteins were secreted into cell culture media for 4 days at 37°C. These media were collected and the secreted receptor proteins were purified. After initial dialysis, multi histidine-tagged receptor proteins were purified with immobilized metal affinity column chromatography and the peak fractions were followed by size exclusion column chromatography as previously described [23]. The purified receptor proteins were dialyzed to storage buffer and they were stored at -80°C until their use for fluorescence polarization peptide binding assay. All purification procedures were performed at 4°C.

### 2.5. Customized synthetic peptides

All peptide fragments used for fluorescence polarization (FP) peptide binding assay were custom-synthesized from Genscript (Piscataway, NJ, USA). Fluorescein isothiocyanate (FITC)-labeled salmon calcitonin (sCT) fragment(22–32) and FITC-labeled AC413(6–25) with Y25P mutation were custom-synthesized from Genscript (Piscataway, NJ, USA). These peptides were used as probes for FP peptide binding assay. Full-length rat amylin (Cat.#H-9475.0500, with HPLC purity above 96%) was purchased from Bachem (Torrance, CA, USA). Pramlintide (Cat.# SML2523-5MG) was purchased from Millipore Sigma (Burlington, MA, USA). Other rat amylin analogs were purchased from Biomatik (Wilmington, Delaware, USA). All peptides used in this study except rat amylin purchased from Bachem and Pramlintide purchased from Millipore Sigma were HPLC-purified with at least 85% purity by Genscript (Piscataway, NJ) or Biomatik (Wilmington, Delaware, USA). Mass spectrometry performed at Genscript or Biomatik confirmed the correct molecular mass of all synthesized peptides. The extinction coefficient of FITC (63,000 M^−1^ ·cm^−1^ at 495 nm, pH 7.0) was used to calculate the concentration of the FITC-labeled peptides. For other peptides, extinction coefficients of tryptophan, tyrosine, and/or cystine residues were used to determine peptide concentrations. The sequences of the peptides used in this study are shown in the Table 1.

**Table 1.**
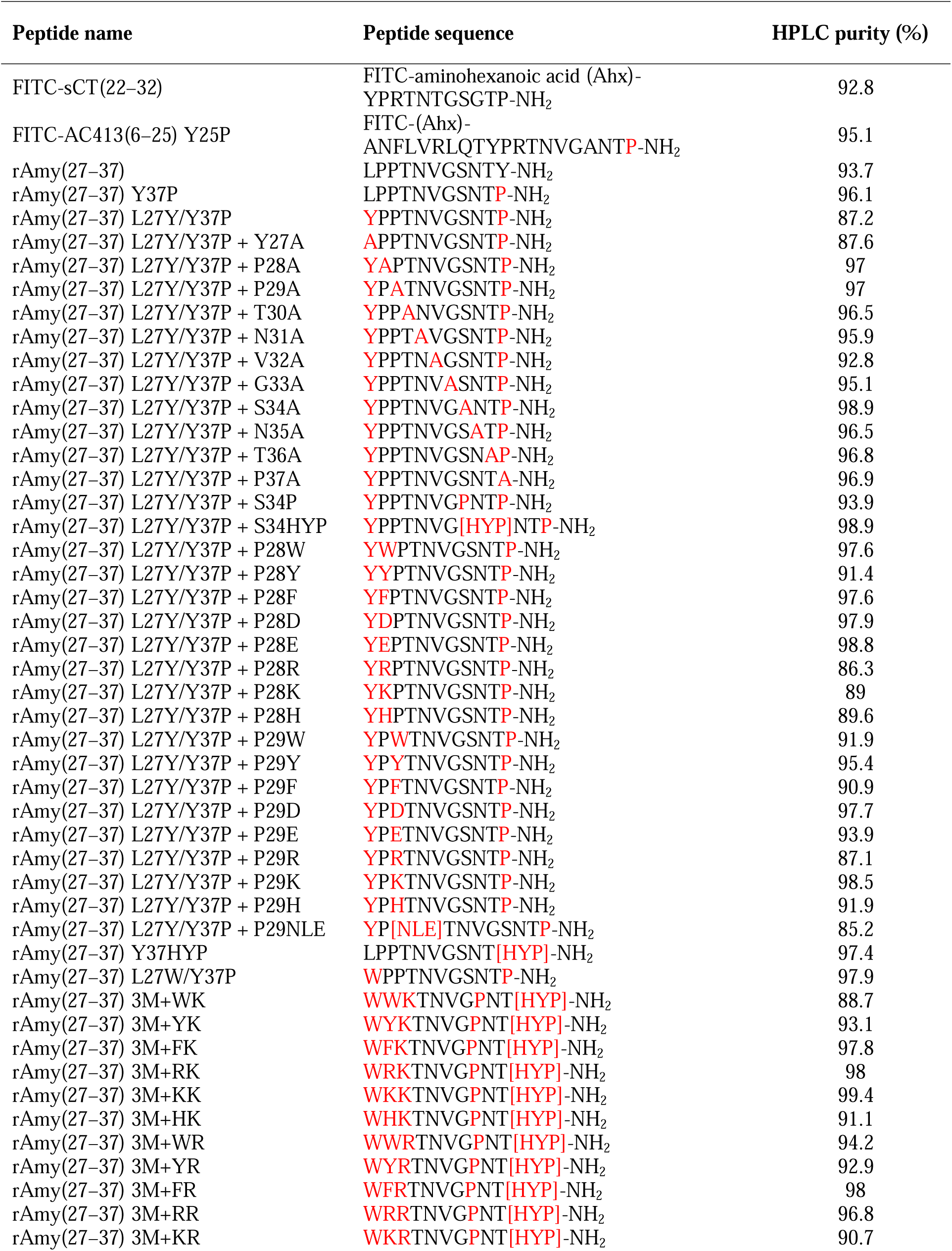
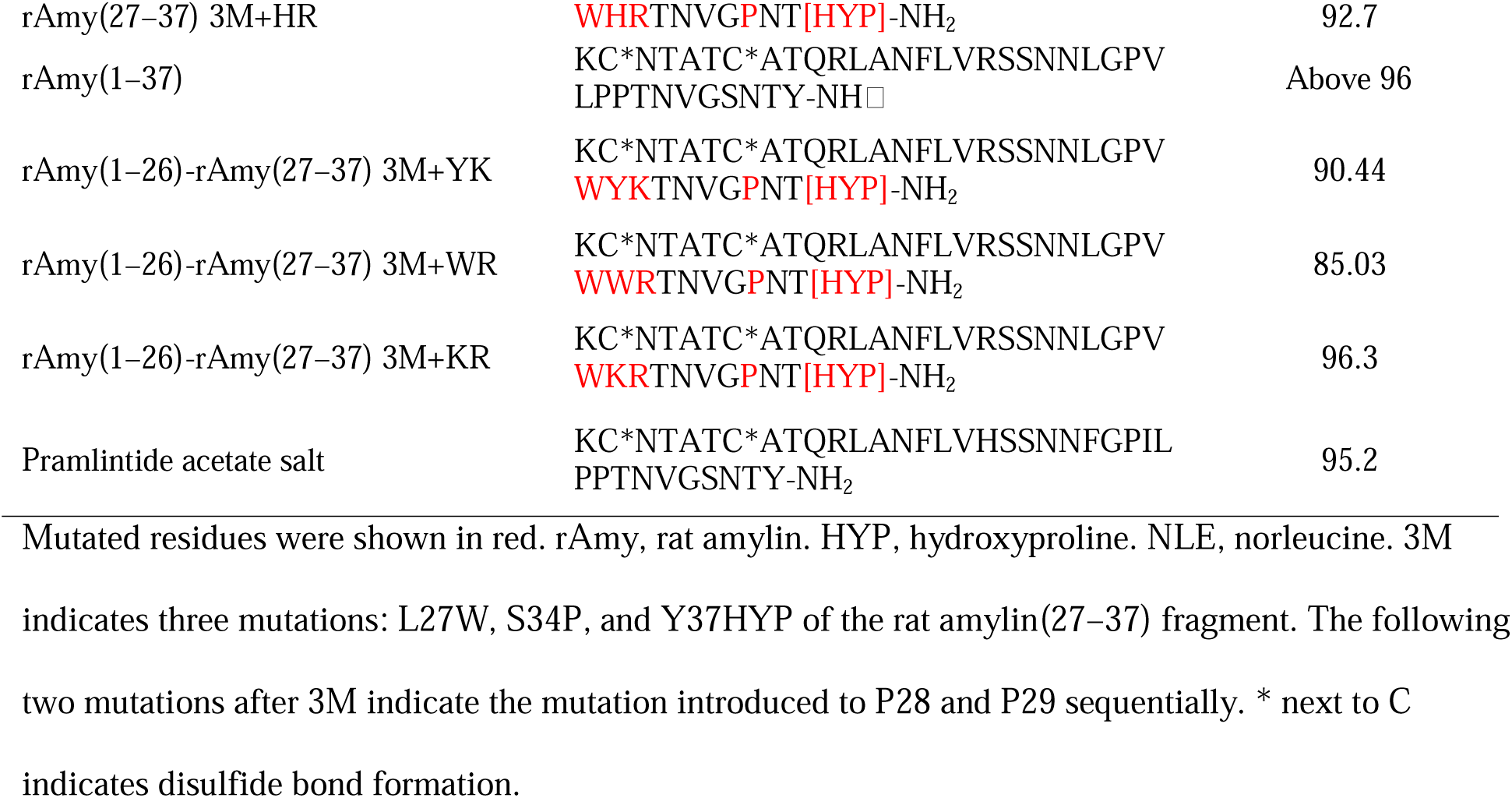
Peptide sequences used in the current study.

### 2.6. Fluorescence polarization/anisotropy (FP) peptide binding assay

The general procedure of FP peptide binding assay with saturation and competition binding formats was previously described [21,24]. 10 nM FITC-labeled sCT(22–32) was used as the peptide probe for CTR ECD binding, while 10 nM FITC-labeled AC413(6-25) with Y25P mutation was used as the peptide probe for all RAMP ECD-CTR ECD fusion proteins. K_D_ values of the FITC-labeled peptide probes used for receptor protein binding were obtained from a saturation binding format. K_D_ of FITC labeled sCT(22–32) for CTR ECD was 500 nM and K_D_ values of FITC-labeled AC413(6–25) Y25P for RAMP1 ECD-CTR ECD, RAMP2 ECD-CTR ECD, and RAMP3 ECD-CTR ECD were 77 nM, 67 nM, and 252 nM. FP binding assay with a competition binding format was established with a fixed concentration (10 nM) of the FITC-labeled peptide probe. The concentration of the purified receptor ECD protein equal to the average K_D_ value was used for the competition binding assay. 500 nM CTR ECD, 77 nM RAMP1 ECD-CTR ECD fusion protein, 67 nM RAMP2 ECD-CTR ECD fusion protein, and 252 nM RAMP3 ECD-CTR ECD fusion protein were used as K_D_ values for competition binding assay. Receptor concentrations equal to the K_D_ value produced approximately 50% of maximum anisotropy. Varying concentrations of a competitive peptide was used for the competition binding assay and the K_I_ value of a competitive peptide was obtained from the competition binding curves. Non-linear regression equations used for saturation and competition binding assay were user-defined for anisotropy calculation as previously described [24,25].

A SpectraiD5 (Molecular Devices, San Jose, CA) was used to evaluate the level of fluorescence polarization/anisotropy. Background signal from the plates was subtracted for accurate anisotropy calculation. For the SpectraiD5, G factor (0.38 for FITC-labeled sCT(22–32), 0.41 for FITC-labeled AC413(6–25) Y25P) was used to correct the instrumental bias for anisotropy calculation. The polarization (mP) of the free FITC-peptide probes without receptor protein binding was set close to 50 mP. When total fluorescence intensity of the FITC-labeled probes was changed with receptor protein interaction by more than 10% of the initial value, the anisotropy was adjusted to reflect the change of total fluorescence intensity as previously described [24]. PRISM 5.0 (GraphPad software, San Diego, CA) was used to produce non-linear regression curves of anisotropy as previously described [21,24]. Two technical replicates were used to obtain anisotropy vales of a testing peptide at each concentration of receptor protein. The mean anisotropy of two replicates was presented in the representative peptide-binding curves. S.E.M. of the anisotropy of the two replicates were presented as error bars. When the error bars were shorter than the height of symbols, they were not shown in the representative curves. At least three independent experiments were performed to obtain at least three K_I_ values of the testing peptide. Mean and standard deviation (S.D.) of the testing peptide K_I_ values were calculated and were shown in the Table 2.

**Table 2.**
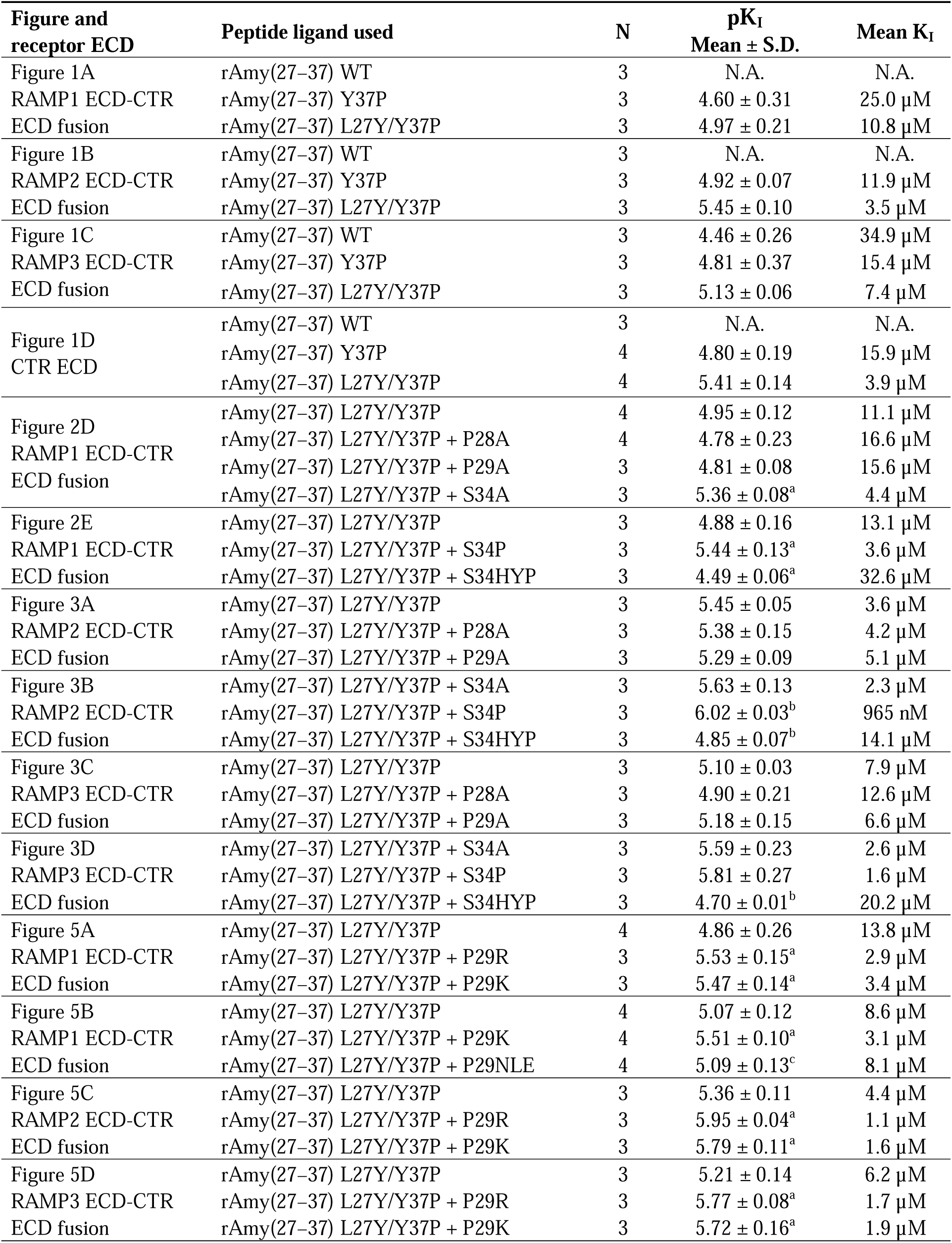
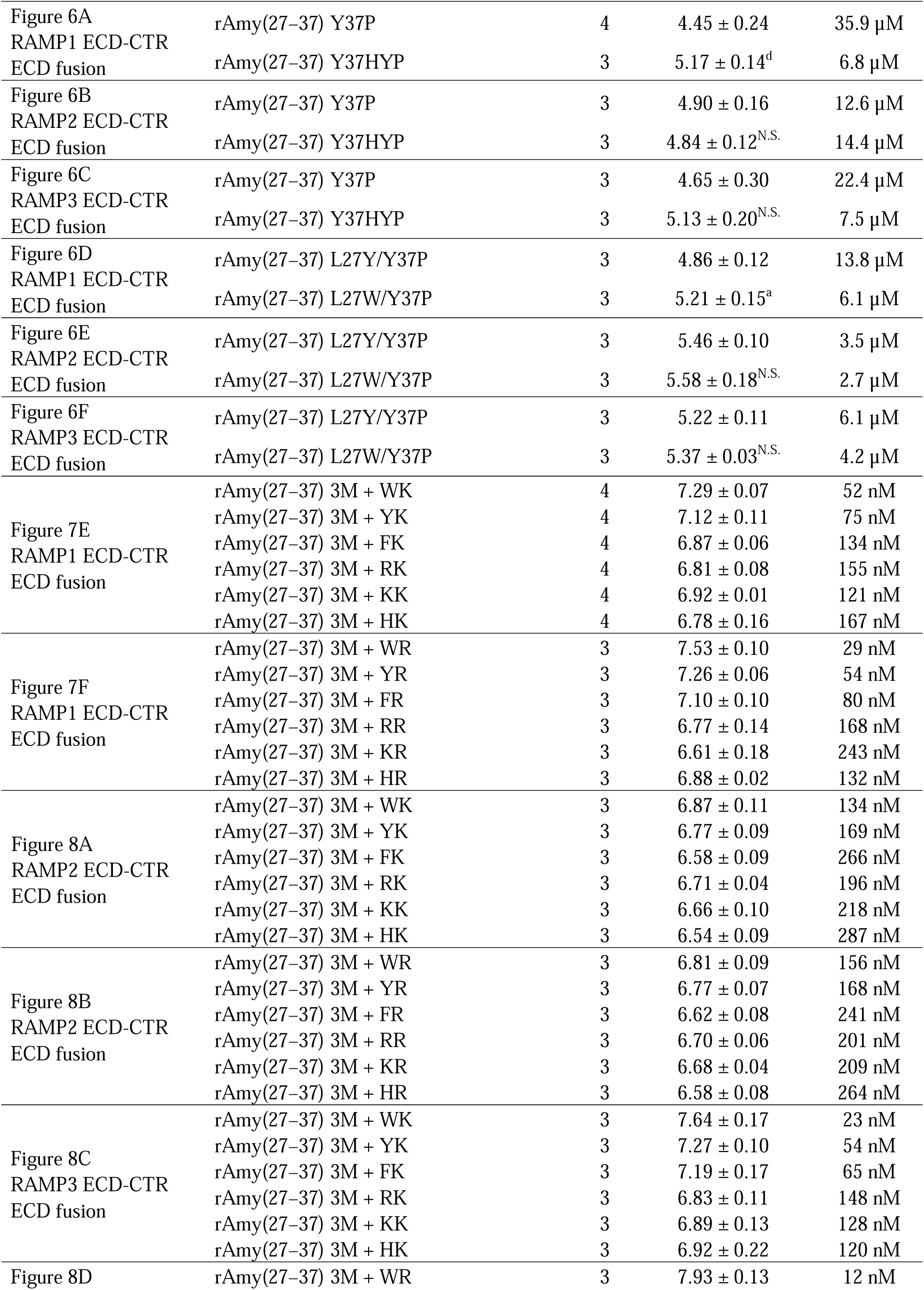
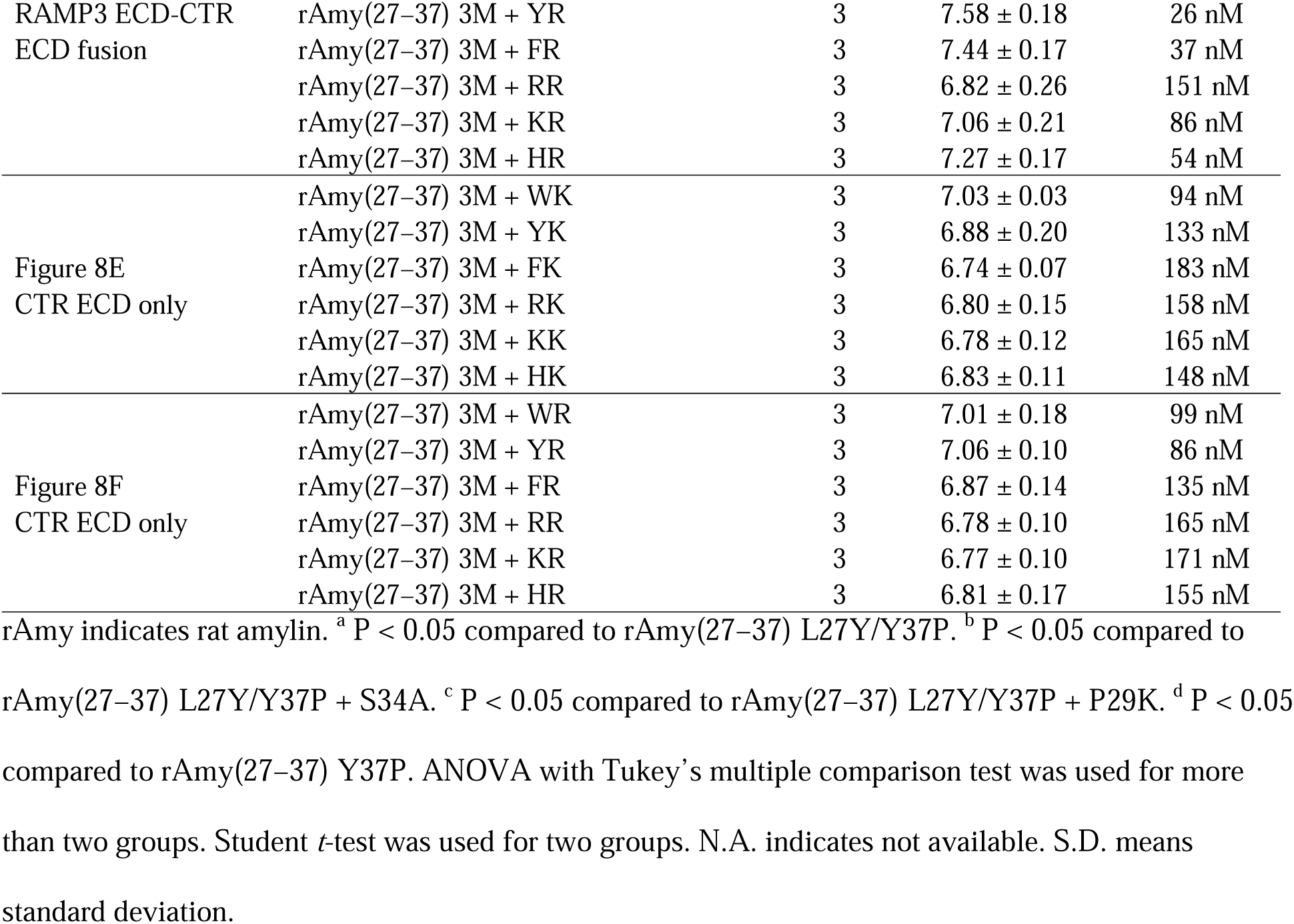
Peptide ligand affinity measured in the present study.

### 2.7. FP competitive binding assay with one fixed concentration of peptides

Alanine-scanning mutagenesis and additional mutations of the rat amylin(27–37) fragment were introduced to investigate mutational effects on peptide ligand binding for calcitonin and amylin receptor ECDs. In Figure 2 and 4, the competitive binding assay format with one fixed peptide concentration was used and the relative competition of the peptides for receptor protein binding was shown as a bar graph. 30 µM peptides were used for RAMP1/3 ECD-CTR ECD fusion proteins (amylin receptor 1/3 ECD), while 10 µM peptides were used for RAMP2 ECD-CTR ECD fusion protein (amylin receptor 2 ECD). These peptide concentrations were determined where the peptide ligand, rAmy(27–37) with L27Y/Y37P mutations approximately decreased the control anisotropy values by half. Compared to the anisotropy value of the peptide ligand, rAmy(27–37) with L27Y/Y37P mutations, higher anisotropy achieved by a mutation suggests a decreased binding (less competition) for receptor proteins, whereas lower anisotropy suggests an increased binding (more competition) for receptor proteins by a mutation. Two technical replicates were used for each sample for one individual experiment. At least three independent experiments were performed to obtain at least six replicates.

### 2.8. Bioluminescence resonance energy transfer (BRET) assay for calcitonin and amylin receptor activation

The general principle and application of using NanoLuc luciferase and Venus-tagged mini G protein for G protein-coupled receptor activation were previously described [18,22]. Amylin receptors are the heterodimer complex of a calcitonin receptor and a receptor activity-modifying protein (RAMP). The human amylin receptor was reconstituted by co-transfecting HEK293T cells with the full-length human calcitonin receptor and one type of three full-length human RAMPs. For transient transfection, HEK293T cells (10^6^ cells/well) were seeded to a six-well plate and were incubated overnight at 37, 5% CO_2_. Next day, HEK293T cells were transfected with the DNA constructs of Venus-tagged mini Gs (2000 µg/well), full-length CTR that was C-terminally tagged with NanoLuc luciferase (100 µg/well), and one type of full-length RAMPs (100 µg/well) for 2 days at 37, 5% CO_2_. The calcitonin receptor activation was evaluated by transfecting HEK293T cells with DNA constructs of Venus-tagged mini Gs (2000 µg/well), full-length CTR that was C-terminally tagged with NanoLuc luciferase (100 µg/well), and pcDNA vector plasmid without any insert (100 µg/well) for 2 days at 37, 5% CO_2_. The same amount of pcDNA vector plasmid as the amount of RAMP DNA plasmid was used for the calcitonin receptor activation to achieve consistent transfection efficiency. Polyethylenimine (PEI) at 1:1.5 ratio (DNA:PEI, w/w) was used as a transfection reagent [19].

After transfection, the HEK293T cells were detached from plate wells by trituration with 5mL of Opti-MEM^®^ I Reduced Serum Medium (Gibco^TM^, Cat.# 31985070) per well. The 5 mL HEK293T cell suspension was combined with the mixture of 10 µL of Nano-Glo^®^ Live Cell Substrate and 90 µL of Nano-Glo^®^ LCS dilution buffer (Promega, Cat.# N2012). The whole mixture of HEK293T with the Nano-Glo^®^ reagents was added to 96 well plates (Greiner Bio-One, Cat.#655075, White, LUMITRAC^TM^, Medium binding) where the testing peptides made by serial dilution were already present. The plates were covered to minimize light exposure and were incubated for 15 min to 20 min at 37 for equilibrium. After incubation, the luminescence signals at 535nm (Lm1) and 485nm (Lm2) were measured with SpectraiD5 (Molecular Devices) using LUM-DUAL setting. The ratio of the luminescence signal measured at 535nm over at 485nm (Lm1/Lm2) was calculated and used as BRET values. The average BRET values from the wells without any peptide were used as a baseline and ΔBRET was calculated by subtracting a baseline from measured BRET values. Three technical replicates were used for each peptide concentration. The mean and S.E.M. of ΔBRET as error bars from the three replicates were presented in the representative ΔBRET curves. When the error bars of the three technical replicates were shorter than the height of the symbol, they were not shown in the representative ΔBRET curves. At least three independent experiments were performed to obtain at least three pEC_50_ values of the testing peptide. Mean and S.D. of the testing peptide pEC_50_ were calculated and were shown in the Table 3.

**Table 3.** Potency for amylin receptor activation obtained from BRET assay. ^a^ P < 0.05 compared to rAmy(1–37). ANOVA with Tukey’s multiple comparison test was used. 3M indicates three mutations of rat amylin(27–37): L27W, S34P, and Y37HYP (hydroxyproline). The following two mutations after 3M indicate the mutation introduced to P28 and P29 sequentially. rAmy, rat amylin. S.D. means standard deviation.

| Activated receptor | Peptide agonist used | N | pEC<sub>50</sub><br>Mean ± SD | Mean<br>EC<sub>50</sub> (nM) |
| --- | --- | --- | --- | --- |
| Figure 9A<br>Amylin receptor 1 | rAmy(1–37) | 4 | 7.04 ± 0.05 | 91 |
|  | rAmy(1–26)-rAmy(27–37) 3M+ YK | 3 | 8.04 ± 0.14<sup>a</sup> | 9 |
|  | rAmy(1–26)-rAmy(27–37) 3M+ WR | 3 | 7.91 ± 0.11<sup>a</sup> | 12 |
|  | rAmy(1–26)-rAmy(27–37) 3M+ KR | 3 | 7.94 ± 0.11<sup>a</sup> | 11 |
|  | Pramlintide | 3 | 7.13 ± 0.14 | 74 |
| Figure 9B<br>Amylin receptor 2 | rAmy(1–37) | 4 | 7.11 ± 0.15 | 77 |
|  | rAmy(1–26)-rAmy(27–37) 3M+ YK | 3 | 8.04 ± 0.06<sup>a</sup> | 9 |
|  | rAmy(1–26)-rAmy(27–37) 3M+ WR | 3 | 7.84 ± 0.24<sup>a</sup> | 14 |
|  | rAmy(1–26)-rAmy(27–37) 3M+ KR | 3 | 7.99 ± 0.17<sup>a</sup> | 10 |
|  | Pramlintide | 3 | 7.13 ± 0.12 | 74 |
| Figure 9C<br>Amylin receptor 3 | rAmy(1–37) | 3 | 7.19 ± 0.06 | 64 |
|  | rAmy(1–26)-rAmy(27–37) 3M+ YK | 3 | 7.13 ± 0.15 | 75 |
|  | rAmy(1–26)-rAmy(27–37) 3M+ WR | 3 | 7.47 ± 0.09 | 34 |
|  | rAmy(1–26)-rAmy(27–37) 3M+ KR | 3 | 7.30 ± 0.19 | 51 |
|  | Pramlintide | 3 | 7.09 ± 0.23 | 82 |
| Figure 9D<br>Calcitonin receptor | rAmy(1–37) | 4 | 7.13 ± 0.09 | 74 |
|  | rAmy(1–26)-rAmy(27–37) 3M+ YK | 3 | 8.18 ± 0.11<sup>a</sup> | 7 |
|  | rAmy(1–26)-rAmy(27–37) 3M+ WR | 3 | 7.84 ± 0.15<sup>a</sup> | 14 |
|  | rAmy(1–26)-rAmy(27–37) 3M+ KR | 3 | 7.97 ± 0.22<sup>a</sup> | 11 |
|  | Pramlintide | 3 | 7.15 ± 0.02 | 72 |

### 2.9. Homology model structure building

SWISS-MODEL (available in Expasy web server) and Pymol (Schrodinger) was used for making the homology models of mutated amylin-bound amylin receptor structures and their representation. The cryo-EM structure of human amylin receptor 1 with rat amylin bound (PDB 7TYF) was used as a template for building homology models for Figure 7B–7D. The amylin receptor with each mutated rat amylin peptide was modeled with SWISS-MODEL. These models were visualized by using Pymol and the mutation of proline to hydroxyproline was made with PyTMs plugin installed in Pymol [26].

### 2.10. Statistical analysis

Student’s *t*-test (two-tailed) was used for the statistical analysis of two groups. One-way ANOVA and following Tukey’s or Dunnet’s *post hoc* test were performed for the comparison of more than two groups. PRISM 5.0 (GraphPad software, San Diego, CA) was used for all statistical analyses. Results with P < 0.05 were considered as a statistically significant difference.

## 3. Results

### 3.1. Design of a backbone rat amylin(27–37) peptide with micromolar affinity for amylin receptor ECDs

The amylin receptor has the extracellular domain (ECD) where a peptide ligand binds presumably at the initial stage of amylin receptor interaction. Amylin receptor ECDs were reconstituted *in vitro* with the fusion proteins of calcitonin receptor ECD and one type of RAMP ECDs with a flexible linker [21,23]. An eleven amino acid salmon calcitonin (sCT) fragment and an antagonistic amylin analog AC413 fragment with/without one mutation (Y25P mutation) have shown micromolar IC_50_ or affinity for these amylin receptor ECDs [20,21,23,24]. However, when a short rat amylin(27–37) fragment was used for FP competition peptide binding assay, it did not show any binding for amylin receptor ECDs at maximal 100 µM concentration (Figure 1). To achieve testable micromolar affinity of the rat amylin(27–37) fragment, two mutations were introduced: Y37P and L27Y mutations. Y37P mutation was previously shown to significantly increase the binding affinity of rat amylin fragment for calcitonin and amylin receptor ECDs [23]. In addition, L27Y mutation came from the sequence of sCT showing micromolar affinity for amylin receptor ECDs. The L27Y and Y37P double mutations of rat amylin(27–37) markedly increased the binding for amylin receptor ECDs with testable micromolar affinity (K_I_ 4 ∼ 11 µM) (Figure 1 and Table 2). These mutations also increased the binding affinity of rat amylin(27–37) for calcitonin receptor ECD similarly to the affinity increases shown for the amylin receptor ECDs (Figure 1D and Table 2) making rat amylin(27–37) with L27Y and Y37P double mutations non-selective for either amylin receptor ECD or calcitonin receptor ECD.

**Figure 1.**
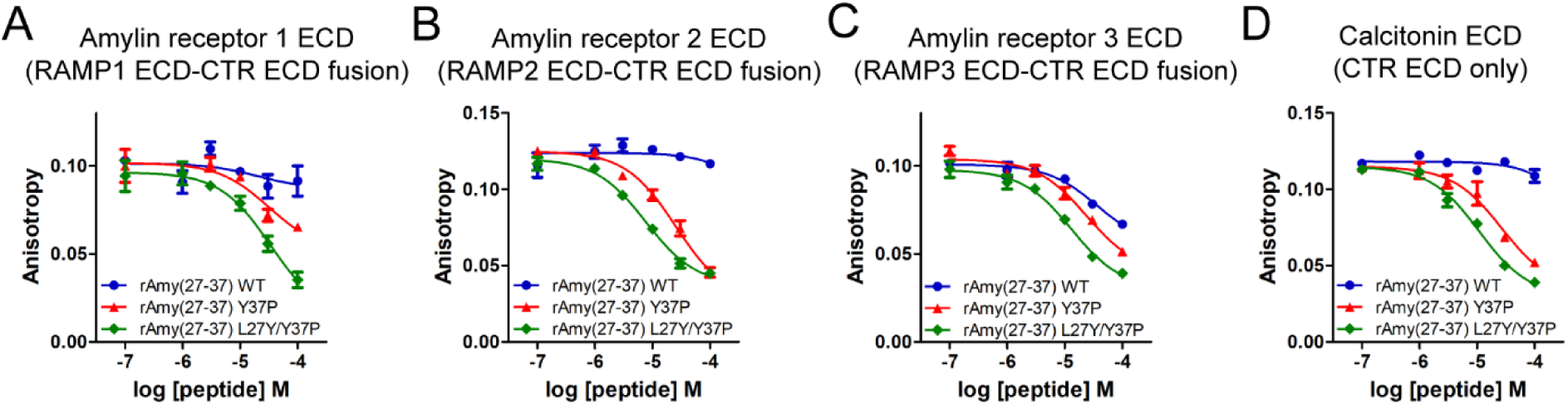
Rat amylin(27–37) peptide binding for amylin receptor ECDs and calcitonin receptor ECD evaluated with fluorescence polarization (FP) peptide binding assay. RAMP1 ECD-CTR ECD and RAMP2 ECD-CTR ECD fusion proteins were produced from HEK293T cells. RAMP3 ECD-CTR ECD fusion protein was produced from HEK293S GnTI^-^ cells. FITC-AC413(6–25) with Y25P mutation was used as the FITC-labeled peptide probe for RAMP1/2/3 ECD-CTR ECD fusion proteins. A, B, and C) Representative peptide binding for amylin receptor 1, 2, and 3 ECD from three independent experiments was shown. D) A backbone rat amylin(27–37) peptide binding for calcitonin receptor (CTR) ECD alone. CTR ECD was produced from HEK293T cells. FITC-sCT(22–32) was used as an FITC-labeled peptide probe for CTR ECD. Representative peptide binding curves were shown from at least three independent experiments.

### 3.2. Alanine-scanning mutagenesis of a backbone peptide rat amylin(27–37) L27Y/Y37P

Alanine-scanning mutagenesis was introduced to the backbone peptide rat amylin(27–37) L27Y/Y37P to investigate key amino acids of the rat amylin fragment to amylin receptor ECD binding. In Figure 2A–2C, Y27A mutation markedly decreased the peptide binding. Alanine mutation of T30, V32, G33 and P37 of the backbone peptide almost abolished the amylin receptor ECD binding suggesting that these residues were critical for binding. Alanine mutation of the backbone peptide N31, N35, and T36 residues moderately decreased the peptide binding. Alanine mutation of rat amylin(27–37) P28 and P29 showed almost no effect on peptide binding for amylin receptor ECDs. Interestingly, S34A mutation increased the peptide binding for amylin receptor ECDs. This result with S34A mutation was consistent with the previous report showing increased sCT(22–32) fragment affinity for calcitonin receptor ECD when the corresponding sCT serine residue (S29) was mutated to alanine [21].

**Figure 2.**
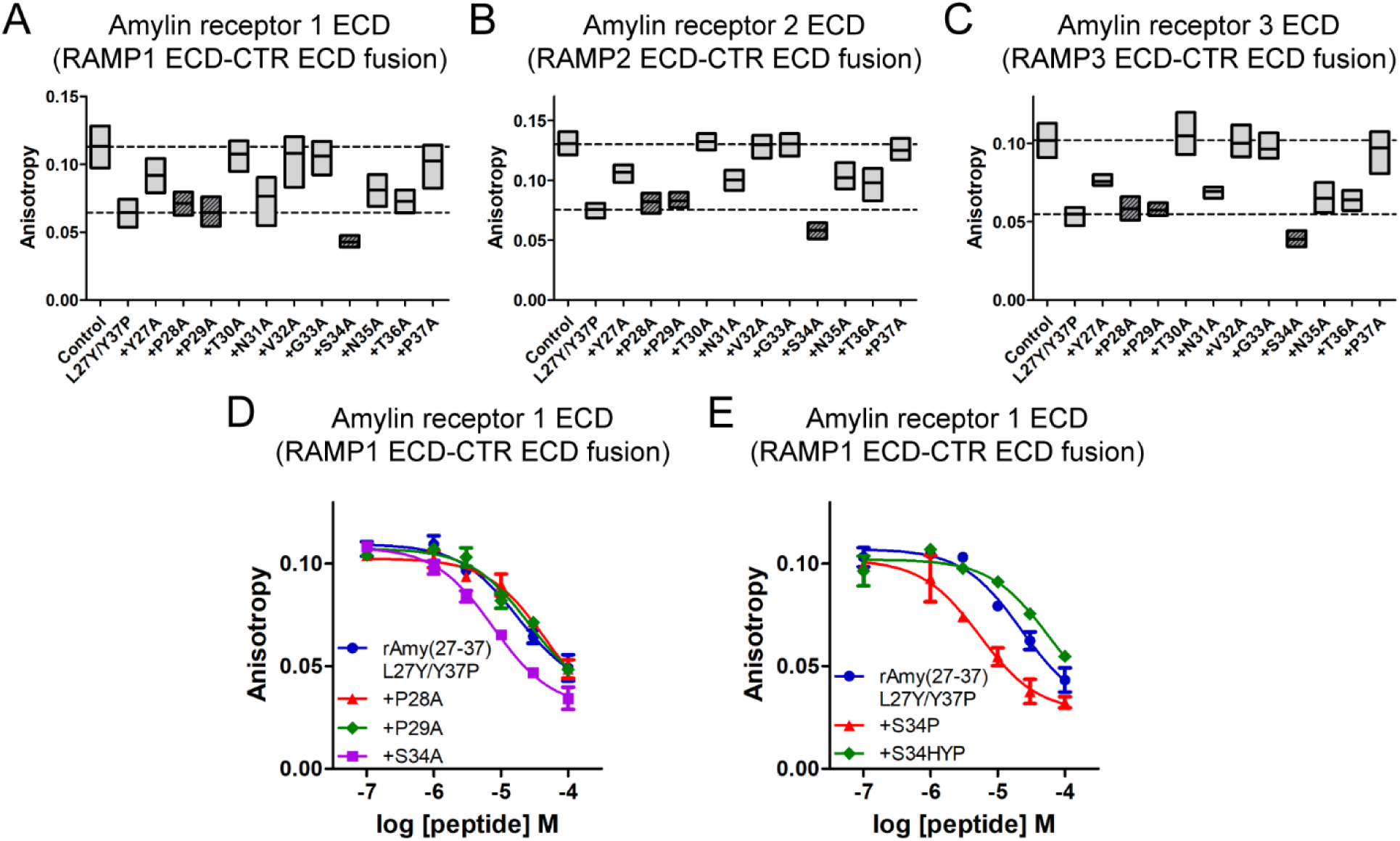
Alanine-scanning mutagenesis of rat amylin(27–37) L27Y/Y37P and the mutational effects on amylin receptor ECD binding. A–C) FP peptide binding assay with one fixed concentration of mutated rat amylin(27–37) for amylin receptor ECDs. 30 μM rat amylin(27–37) was used for amylin receptor 1/3 ECD binding, while 10 μM rat amylin(27–37) was used for amylin receptor 2 ECD binding. AC413(6–25) with Y25P mutation was used as an FITC-labeled peptide probe for the FP peptide binding assay. For comparison, anisotropy values of Control (no competitive peptide) and of rat amylin(27–37) L27Y/Y37P were indicated with dotted lines. Two technical replicates were used for each experiment. Six anisotropy values from three independent experiments were combined and the average anisotropy was shown with the box plot indicating the highest and the lowest values and the mean. P28A, P29A and S34A that were further investigated in Figure 3 were shown in dark gray and with tilted lines. D) FP peptide binding assay with rat amylin(27–37) L27Y/Y37P plus P28A, P29A, or S34A mutation. E) FP peptide binding assay with rat amylin(27–37) L27Y/Y37P plus S34P or S34HYP (hydroxyproline) mutation. AC413(6–25) with Y25P mutation was used as an FITC-labeled peptide probe for the FP peptide binding assay. Representative peptide binding curves were shown from at least three independent experiments.

Mutational effects of the backbone peptide P28, P29, and S34 residues were further investigated with the RAMP1 ECD-CTR ECD fusion protein in full competition binding assay. As expected, P28A and P29A mutations did not show a significant change in pK_I_ affinity values for amylin receptor 1 ECD binding, while S34A mutation significantly increased the backbone peptide affinity by 2.5-fold (Figure 2D and Table 2). Rat amylin S34 corresponds to sCT S29 residue when the sequence was aligned [23]. Since the corresponding sCT S29P mutation significantly increased the peptide binding affinity for calcitonin receptor ECDs by 5-fold [21], the mutation to proline was also introduced to S34 of the rat amylin backbone peptide. Consistently, S34P mutation of the rat amylin backbone peptide significantly increased peptide affinity for amylin receptor 1 ECD binding by 3.6-fold (Figure 2E and Table 2). S34HYP mutation where the additional hydroxyl group of hydroxyproline (HYP) deemed to generate steric clash with the peptide binding pocket of calcitonin receptor ECD. S34HYP mutation markedly decreased the peptide affinity for amylin receptor 1 ECD by 9-fold, when compared to the peptide affinity with S34P mutation (Figure 2E and Table 2). These results suggest that relatively small side chains of alanine or proline at position 34 of the backbone peptide have a better fit for amylin receptor 1 ECD binding than the relatively bulky side chain of hydroxyproline.

To make sure that the mutational effects of P28A, P29A, S34A, S34P, and S34HYP were also observed with amylin receptor 2/3 ECD, FP peptide binding assay was performed with amylin receptor 2/3 ECD. Consistently, these mutational effects of the backbone peptide P28, P29 and S34 residues were conserved with amylin receptor 2/3 receptor ECD binding (Figure 3 and Table 2).

**Figure 3.**
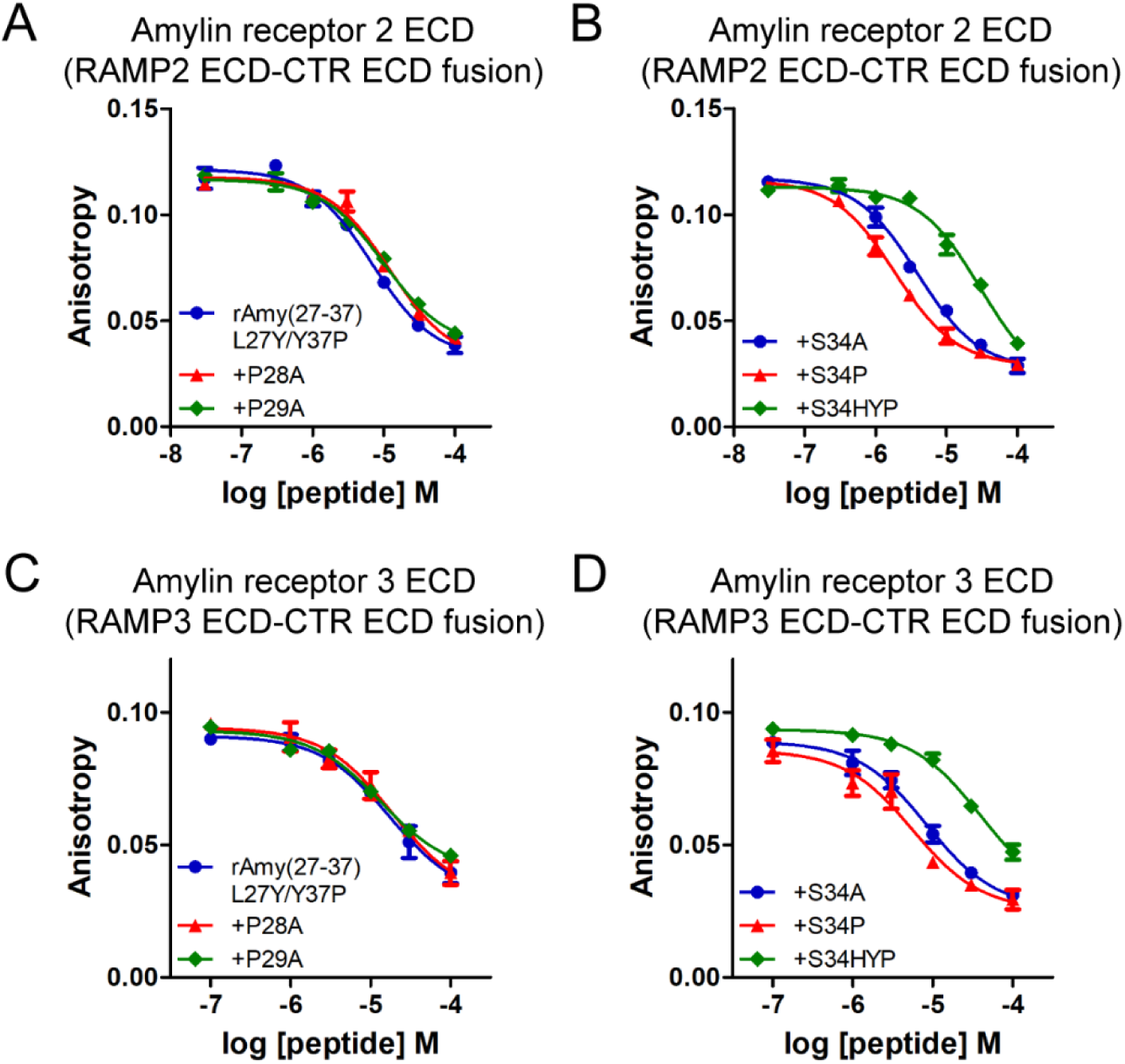
Full competition binding assay with P28, P29 and S34 mutations of rat amylin(27–37) L27Y/Y37P for amylin receptor 2/3 ECD binding. A–D) FP peptide binding assay with rat amylin(27–37) L27Y/Y37P plus P28A, P29A, S34A, S34P, or S34HYP (hydroxyproline) mutation for amylin receptor 2/3 ECD. AC413(6–25) with Y25P mutation was used as an FITC-labeled peptide probe for the FP peptide binding assay. Representative peptide binding curves were shown from three independent experiments.

### 3.3. More mutations of rat amylin backbone peptide P28 and P29 residues

P28 and P29 residues of the backbone peptide were further explored with more mutations since the side chain of these two prolines appeared dispensable for amylin receptor ECD binding. Structural studies including sCT and calcitonin receptor ECD suggest that P28 and P29 were relatively distant from the peptide binding pocket of the calcitonin and amylin receptor ECDs and that adding bulky residues would not create steric hindrance for peptide binding [15,27]. Mutations to bulky residues (tryptophan, tyrosine, and phenylalanine) or to charged residues (aspartate, glutamate, arginine, lysine, and histidine) were introduced to either P28 or P29. Except P28D mutation, other mutations at P28 showed minor change in peptide binding for amylin receptor 1 ECD (Figure 4A). When amylin receptor 2/3 ECDs were used for the binding assay, the pattern of P28 mutational effects was similar to that for amylin receptor 1 ECD (Figure 4B and 4C). Interestingly, the mutations of P29 to positively charged residues (P29R and P29K mutations) increased the peptide binding for amylin receptor 1 ECD (Figure 4D). In contrast, the mutations of P29 to negatively charged residues (P29D and P29E mutations) markedly decreased the peptide binding (Figure 4D). These results suggest that the positive charge of P29R and P29K mutations is potentially involved in affinity enhancement. Amylin receptor 2/3 ECDs also showed the similar pattern of P29 mutational effects to the results with amylin receptor 1 ECD (Figure 4E and 4F).

**Figure 4.**
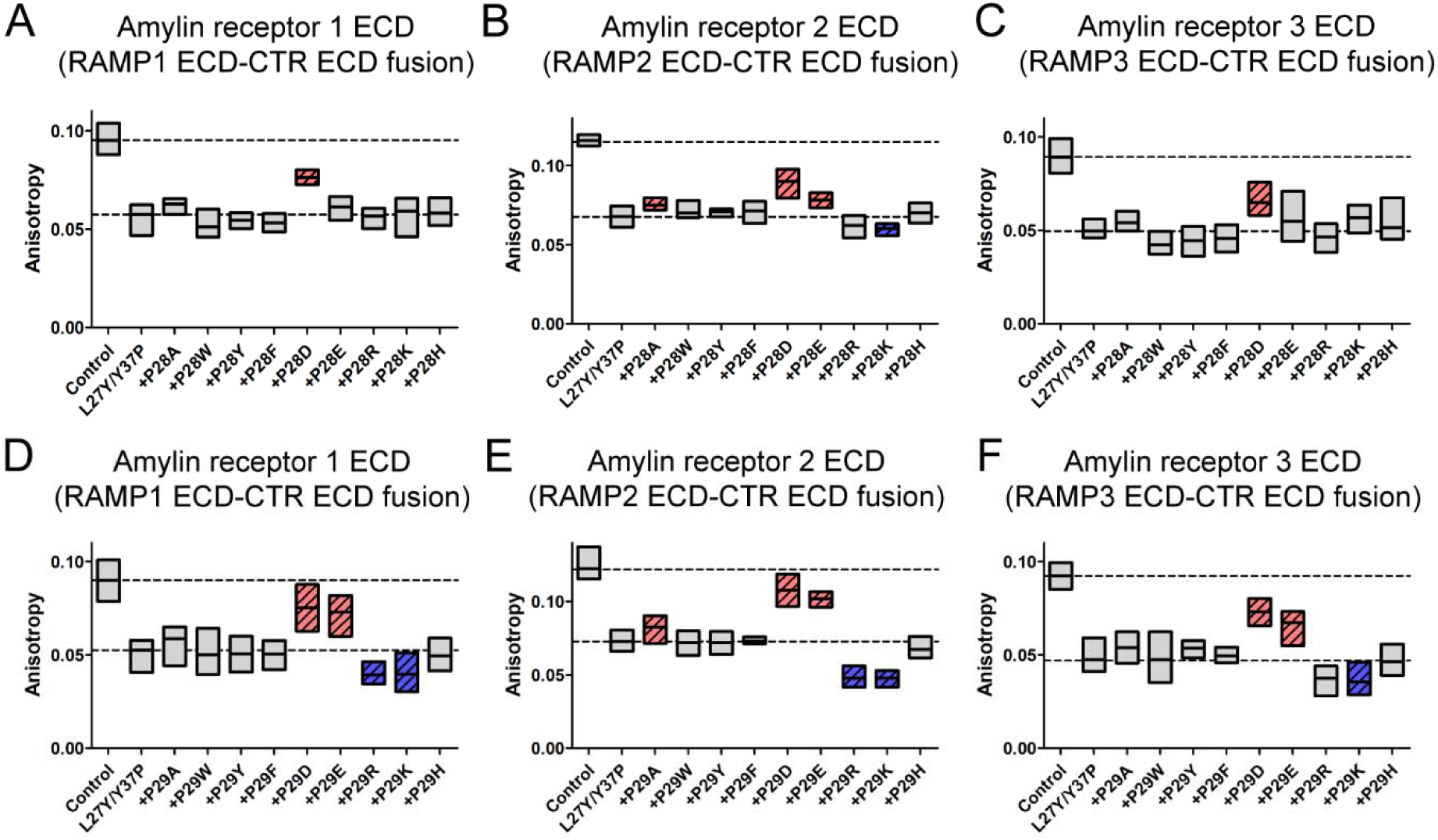
Additional mutations of P28 and P29 of rat amylin(27–37) L27Y/Y37P and their effects on amylin receptor 1/2/3 ECD binding A–C) FP peptide binding assay with rat amylin(27–37) L27Y/Y37P plus P28 mutations. D–F) FP peptide binding assay with rat amylin(27–37) L27Y/Y37P plus P29 mutations. 30 μM rat amylin(27–37) L27Y/Y37P plus P28 and P29 mutations was used for amylin receptor 1/3 ECD, while 10 μM rat amylin(27–37) L27Y/Y37P plus P28 and P29 mutations was used for amylin receptor 2 ECD. AC413(6–25) with Y25P mutation was used as an FITC-labeled peptide probe for the FP peptide binding assay. P28D, P28E, P29A, P29D, and P29E mutations with a statistically significant anisotropy increase were indicated in red and with tilted lines, while P29R and P29K mutations with a statistically significant anisotropy decrease were indicated in blue and with tilted lines. For comparison, anisotropy values of Control (no competitive peptide) and rat amylin(27–37) L27Y/Y37P were indicated with dotted lines. Two technical replicates were used for each experiment. Six anisotropy values from three independent experiments were combined and the average anisotropy was shown with the box plot indicating the highest and the lowest values and the mean. One-way ANOVA with Dunnet’s *post hoc* test was used for statistical analysis. The L27Y/Y37P sample was used as a statistical control for the comparison in Dunnet’s *post hoc* test. P< 0.05 was used for statistical significance.

The mutational effects of P29R or P29K were further investigated with full competition binding assay. P29R or P29K mutation increased the backbone peptide affinity for amylin receptor 1 ECD by 4-fold (Figure 5A and Table 2). To confirm the role of the positive charge in peptide binding affinity, norleucine was introduced to P29 instead of lysine. Norleucine has the same length of the side chain as lysine does. However, it has -CH_3_ at the end of the side chain instead of a positively charged -NH_3_ moiety in lysine such that the effect of the positive charge of lysine is removed. When norleucine was introduced to the backbone peptide P29, the increased affinity produced by P29K mutation disappeared mostly (Figure 5B). This suggests that adding a positive charge to P29 through P29K/R mutation significantly contributes to the increased peptide affinity for amylin receptor 1 ECD. The mutational effects of P29K/R mutation on amylin receptor 2/3 ECD binding were similar to those on amylin receptor 1 ECD binding (Figure 5C, 5D and Table 2).

**Figure 5.**
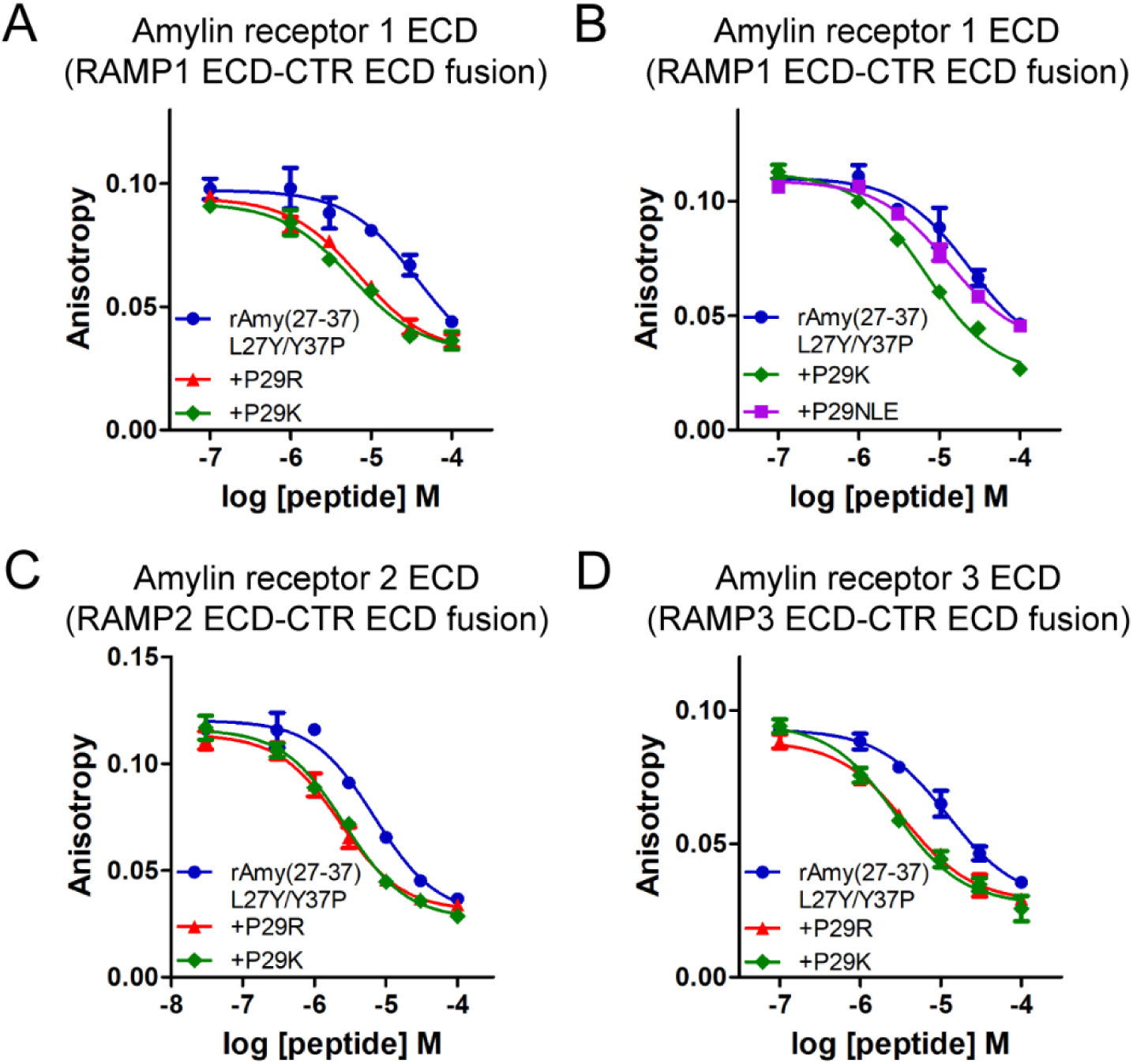
P29R or P29K mutational effects for amylin receptor 1/2/3 ECD binding A) FP peptide binding assay with rat amylin(27–37) L27Y/Y37P plus P29R or P29K mutation and amylin receptor 1 ECD. B) FP peptide binding assay with rat amylin(27–37) L27Y/Y37P plus P29NLE (norleucine) mutation and amylin receptor 1 ECD. C, D) FP peptide binding assay with rat amylin(27–37) L27Y/Y37P plus P29R or P29K mutation for amylin receptor 2/3 ECD. AC413(6–25) with Y25P mutation was used as an FITC-labeled peptide probe for the FP peptide binding assay. Representative peptide binding curves were shown from at least three independent experiments.

### 3.4. Additional mutations that enhanced the affinity of the rat amylin(27–37) fragment

Two more mutations were found to increase the affinity of rat amylin(27–37) for amylin receptor 1 ECD: Y37HYP and L27W mutations. P32HYP mutation of sCT(22–32) fragment was reported to moderately increase peptide affinity for calcitonin and amylin receptor ECDs [21]. When HYP mutation was introduced to Y37 of rat amylin(27–37), the affinity for amylin receptor 1 ECD was further increased by 5-fold compared to the affinity increase by Y37P mutation (Figure 6A and Table 2). Interestingly, Y37HYP mutation did not make significant changes in rat amylin(27–37) affinity for amylin receptor 2 ECD compared to Y37P mutation (Figure 6B and Table 2). In contrast, Y37HYP mutation significantly increased rat amylin(27–37) affinity for amylin receptor 3 ECD by 3-fold compared to Y37P mutation (Figure 6C and Table 2).

**Figure 6.**
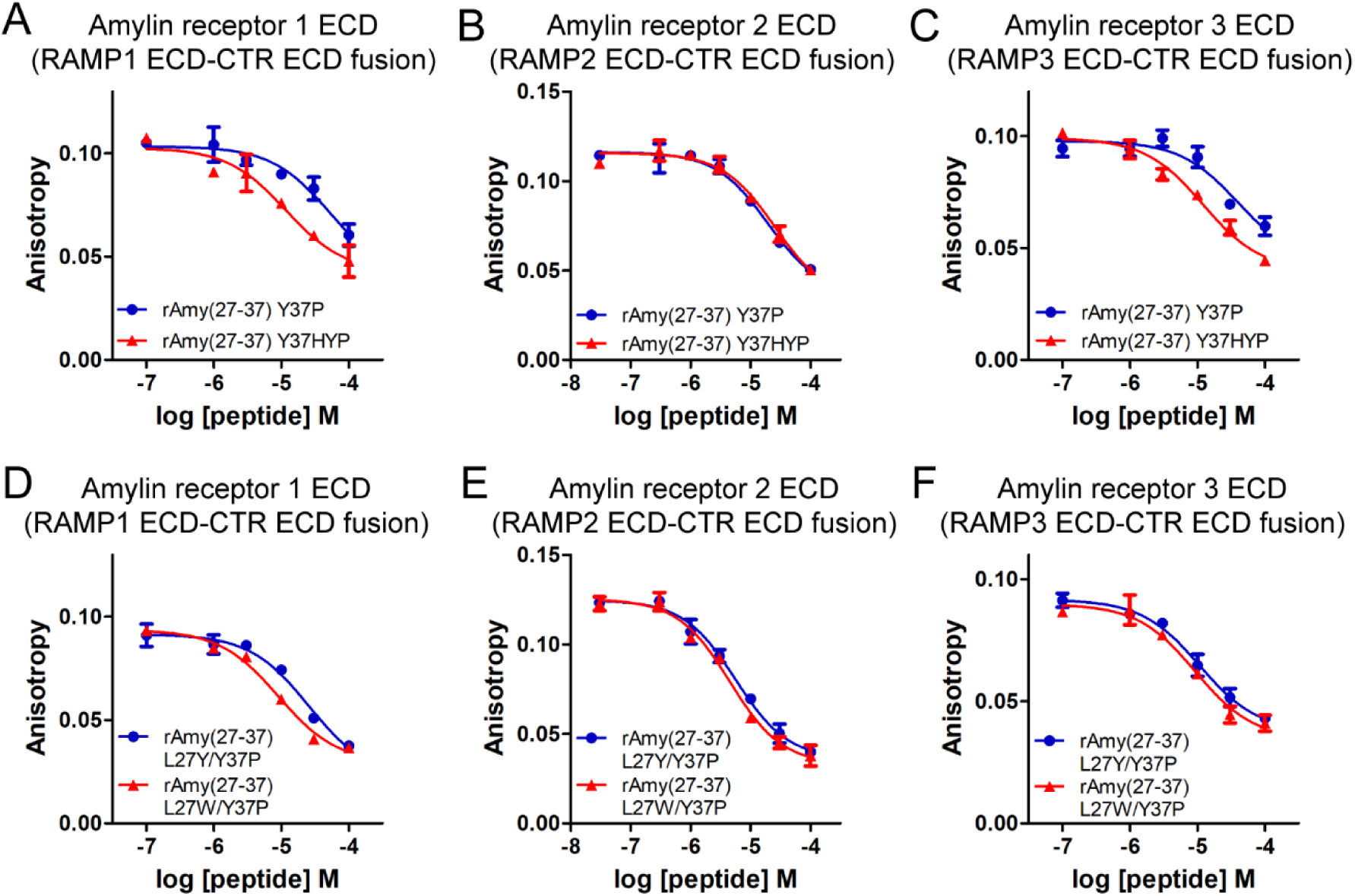
Y37HYP and L27W mutations enhanced peptide binding affinity for amylin receptor 1 ECD. A–C) FP peptide binding assay with rat amylin(27–37) Y37P or Y37HYP mutation and amylin receptor 1/2/3 ECD. D–F) FP peptide binding assay with rat amylin(27–37) L27W/Y37P mutations and amylin receptor 1/2/3 ECD. AC413(6–25) with Y25P mutation was used as an FITC-labeled peptide probe for the FP peptide binding assay. Representative peptide binding curves were shown from at least three independent experiments.

Since L27Y mutation increased rat amylin(27–37) affinity (Figure 1), the effect of L27 mutation to tryptophan that has a bit bulkier side chain than tyrosine was examined. L27W mtation of rat amylin(27–37) was found to increase peptide affinity for amylin receptor 1 ECD by 2-fold compared to L27Y mutation (Figure 6D and Table 2). L27W mutation did not further increase the affinity of rat amylin(27–37) for amylin receptor 2/3 ECD compared to L27Y mutation (Figure 6E, 6F and Table 2).

### 3.5. Design of improved rat amylin(27–37) mutated analogs and their potential interaction mechanism at amylin receptor 1 ECD

Based on affinity-enhancing mutations for amylin receptor ECDs, twelve rat amylin(27–37) mutated analogs were designed by combining all mutations to achieve maximal affinity enhancement (Figure 7A). The structural homology models of these rat amylin(27–37) mutated analogs bound for amylin receptor 1 ECD were shown in Figure 7B–7D. L27W mutation of the rat amylin(27–37) fragment added a bulky tryptophan to L27 position and the tryptophan located in proximity of CTR P100 and rat amylin N21 residues (Figure 7C). Rat amylin P28 was placed outward from the peptide binding pocket such that any tested mutations at P28 did not have the steric clash with CTR (Figure 7B–7D). Rat amylin P29K mutation appeared to be in proximity (3.5) of CTR N135 suggesting that the positive charge of lysine participates in the potential hydrogen bond formation with CTR N135 (Figure 7D). The binding results with P29 to norleucine mutation (Figure 5B) was consistent with the potential interaction of lysine and arginine (P29K/R) with CTR N135 suggested in Figure 7D. Consistently with a previous report [21], rat amylin S34P mutation appeared to fit the peptide binding pocket of CTR more suitably with CTR E123 and N124 (Figure 7C). Rat amylin Y37HYP mutation also appeared to increase the interaction with CTR W79 (Figure 7C).

**Figure 7.**
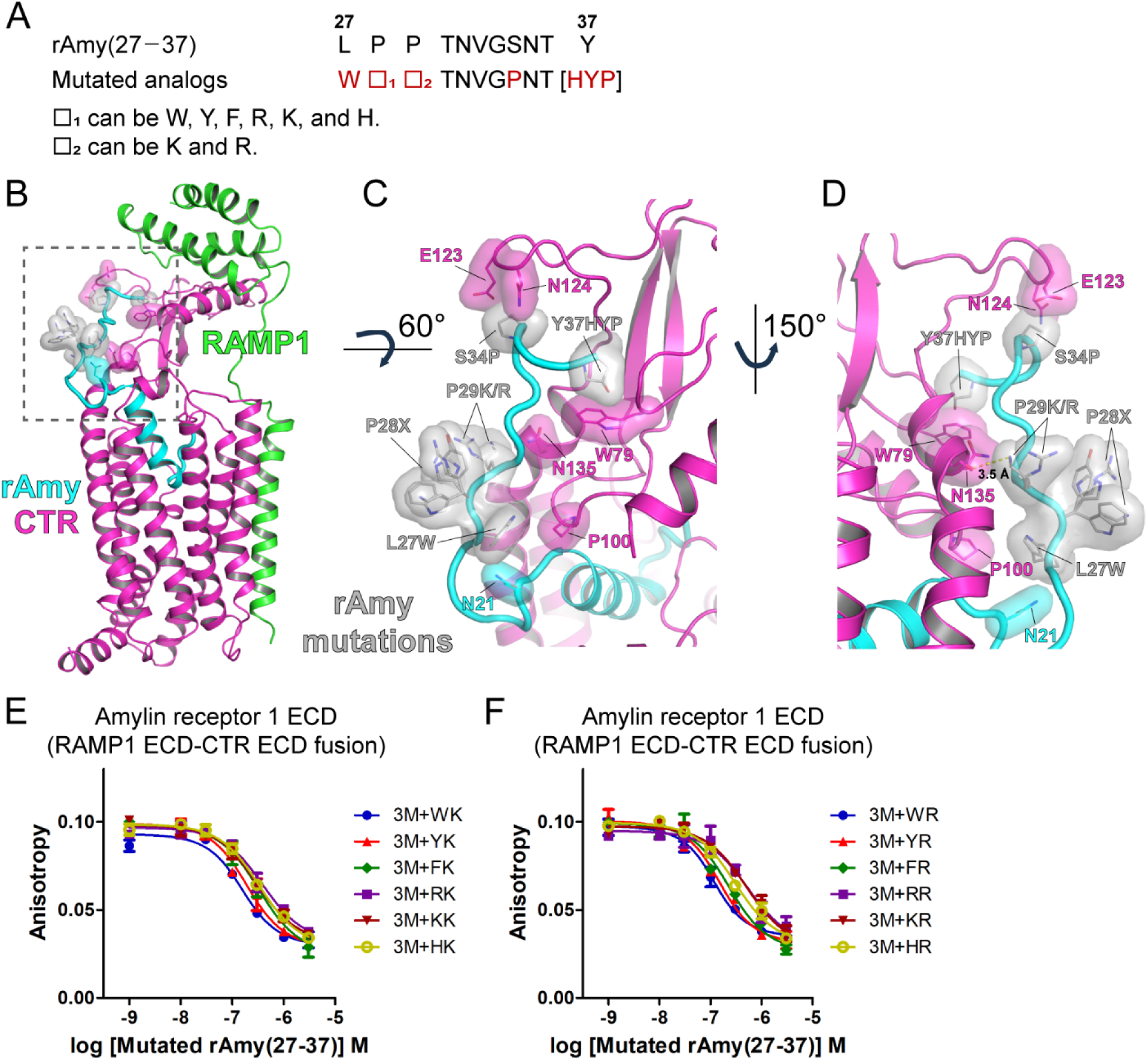
Design of rat amylin(27–37) fragments with enhanced affinity for amylin receptor ECDs. A) Mutated analogs of rat amylin(27–37) with affinity enhancement for amylin receptor ECDs. Mutated sites were shown in red. Rat amylin(27–37) was shown for comparison. B) The structural homology models of mutated analogs of rat amylin(27–37) bound at amylin receptor 1. The cryo-EM structure of human amylin receptor 1 and rat amylin (PDB: 7TYF) was used. Mutated rat amylin residues were shown in gray and with stick and surface representation. The calcitonin receptor residues potentially interacting with the mutated rat amylin residues were shown in purple and with stick and surface representation. Rat amylin N21 in proximity of rat amylin L27W residue was shown in cyan and with stick and surface representation. C and D) Magnified views of the area with the dotted lines in panel B. RAMP1 was hidden for visual clarity. The distance between the main chain of CTR N135 and rAmy(27–37) P29K was 3.5 and was shown as a yellow dotted line. E) FP peptide binding assay with rat amylin(27–37) 3M + WK, YK, FK, RK, KK, and HK and amylin receptor 1 ECD. F) FP peptide binding assay with rat amylin(27–37) 3M + WR, YR, FR, RR, KR, and HR and amylin receptor 1 ECD. 3M indicates three mutations of rat amylin(27–37): L27W, S34P, and Y37HYP (hydroxyproline). The following two mutations after 3M indicate the mutation introduced to P28 and P29 sequentially. AC413(6–25) with Y25P mutation was used as an FITC-labeled peptide probe for the FP peptide binding assay. Representative peptide binding curves were shown from at least three independent experiments. X of P28X means W, Y, F, R, K, and H. CTR, calcitonin receptor; RAMP1, receptor activity-modifying protein 1; rAmy, rat amylin; HYP, hydroxyproline.

The combined mutational effects were examined for amylin receptor ECD binding. A rat amylin(27–37) fragment with L27W, S34P, and Y37HYP mutations was used as an initial rat amylin(27–37) fragment labeled as rAmy(27–37) 3M. On top of that, P28 to W, Y, F, R, K, or H mutation and P29 to K or R mutation were introduced. The additional mutations introduced to P28 and P29 position were labeled sequentially after 3M. The affinity of the rat amylin(27–37) fragment with these mutations for amylin receptor 1 ECD binding ranged from pK_I_ 6.61 to 7.53 (K_I_ 29 nM to 243 nM) (Figure 7E, 7F and Table 2). These affinity values were 103- to 862-fold higher than the affinity of rat amylin(27–37) with Y37P mutation.

For amylin receptor 2/3 ECDs, these combined mutations of rat amylin(27–37) also showed similar affinity enhancement for amylin receptor 2 ECD with pK_I_ 6.54 to 6.87 (K_I_ 134 nM to 287 nM) and for amylin receptor 3 ECD with pK_I_ 6.82 to 7.93 (K_I_ 12 nM to 151 nM), respectively (Figure 8A–8D and Table 2). These mutations showed similar affinity enhancement for CTR ECD alone with pK_I_ 6.74 to 7.06 (K_I_ 86 nM to 183 nM) (Figure 8E, 8F and Table 2). These suggest that the mutated rat amylin(27–37) fragments are not selective for amylin receptor ECDs.

**Figure 8.**
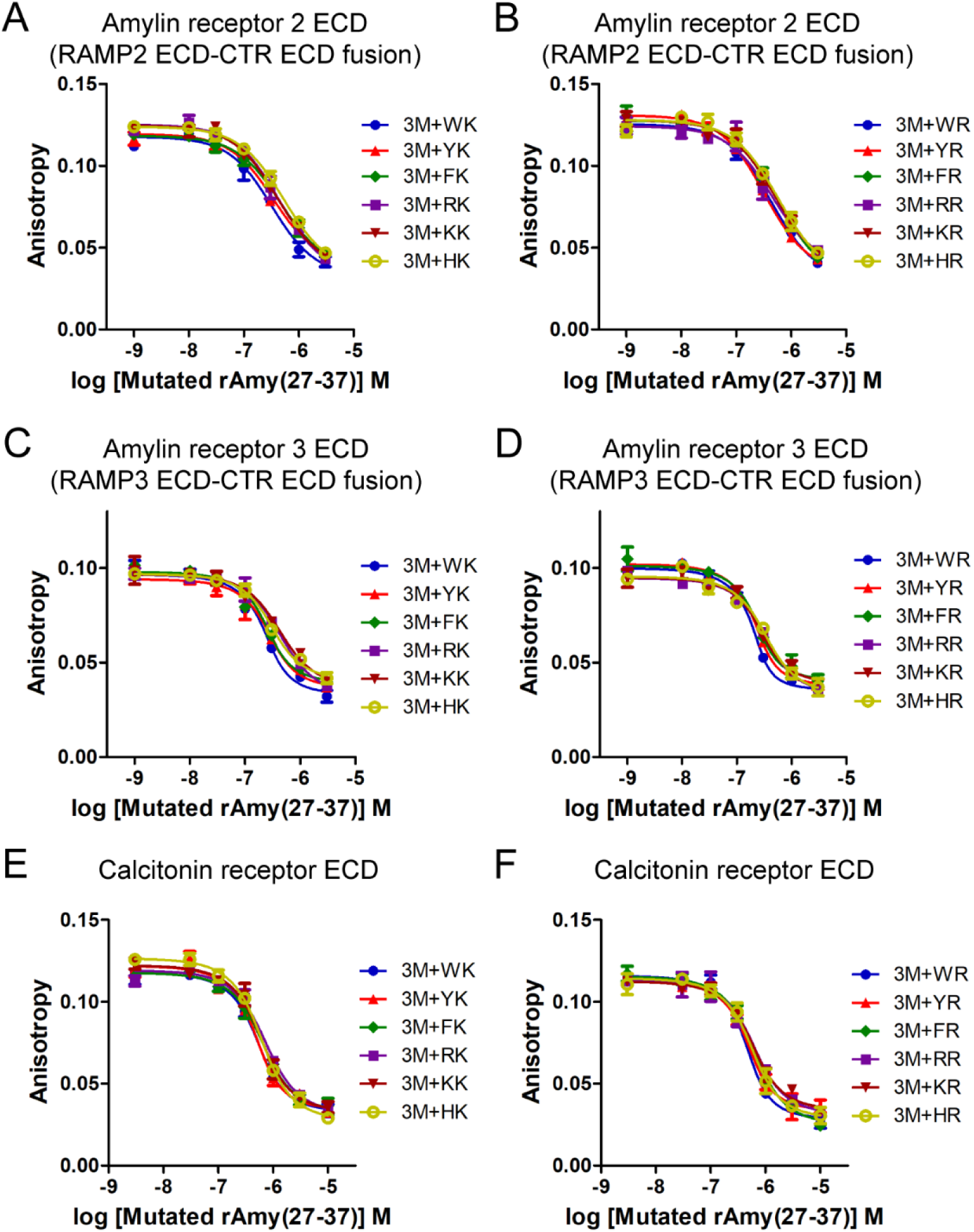
Rat amylin(27–37) fragments with advanced affinity for amylin receptor 2/3 ECDs and CTR ECD. A and B) FP peptide binding assay with rat amylin(27–37) 3M + WK, YK, FK, RK, KK, HK, WR, YR, FR, RR, KR, and HR for amylin receptor 2 ECD. C and D) FP peptide binding assay with rat amylin(27–37) 3M + WK, YK, FK, RK, KK, HK, WR, YR, FR, RR, KR, and HR for amylin receptor 3 ECD. 3M indicates three mutations of rat amylin(27–37): L27W, S34P, and Y37HYP (hydroxyproline). AC413(6–25) with Y25P mutation was used as an FITC-labeled peptide probe for the FP peptide binding assay. E and F) FP peptide binding assay with rat amylin(27–37) 3M + WK, YK, FK, RK, KK, HK, WR, YR, FR, RR, KR, and HR for CTR ECD alone. FITC-sCT(22–32) was used as an FITC-labeled peptide probe for the FP peptide binding assay. The following two mutations after 3M indicate the mutation introduced to P28 and P29 sequentially. Representative peptide binding curves were shown from three independent experiments.

### 3.6. Full-length rat amylin analogs with enhanced potency for amylin receptor activation

Rat amylin(27–37) fragments with affinity-enhancing mutations were used to produce full-length rat amylin analogs. Rat amylin(27–37) with L27W, S34P, and Y37HYP mutations was labeled as rat amylin(27–37) 3M and the additional mutations at P28 and P29 position were labeled sequentially afterward. Three mutated rat amylin(27–37) fragments were selected for making full-length rat amylin analogs: rat amylin(27–37) 3M + YK, rat amylin(27–37) 3M + WR, and rat amylin(27–37) 3M + KR. When these affinity-enhancing mutations were added to full-length rat amylin, all three mutated rat amylin analogs showed enhanced potency for amylin receptor 1/2 activation by 5- to 10-fold compared to endogenous rat amylin and a clinical drug pramlintide (Figure 9A, 9B and Table 3). In contrast, rat amylin potency for amylin receptor 3 activation was not significantly enhanced by these mutations (Figure 9C and Table 3). The potency increases for amylin receptor 3 activation with these mutations were less than 2.5-fold. Rat amylin analogs with these affinity-enhancing mutations also significantly enhanced potency for calcitonin receptor activation by 5- to 11-fold compared to endogenous rat amylin and a clinical drug pramlintide (Figure 9D and Table 3). Calcitonin and amylin activation potency of an approved anti-diabetic drug pramlintide was similar to the potency of rat amylin; pramlintide pEC_50_ was 7.09 to 7.13 (EC_50_ 74 to 82 nM) for amylin receptor 1–3 activation and pramlintide pEC_50_ was 7.15 (EC_50_ 72 nM) for calcitonin receptor activation (Table 3).

**Figure 9.**
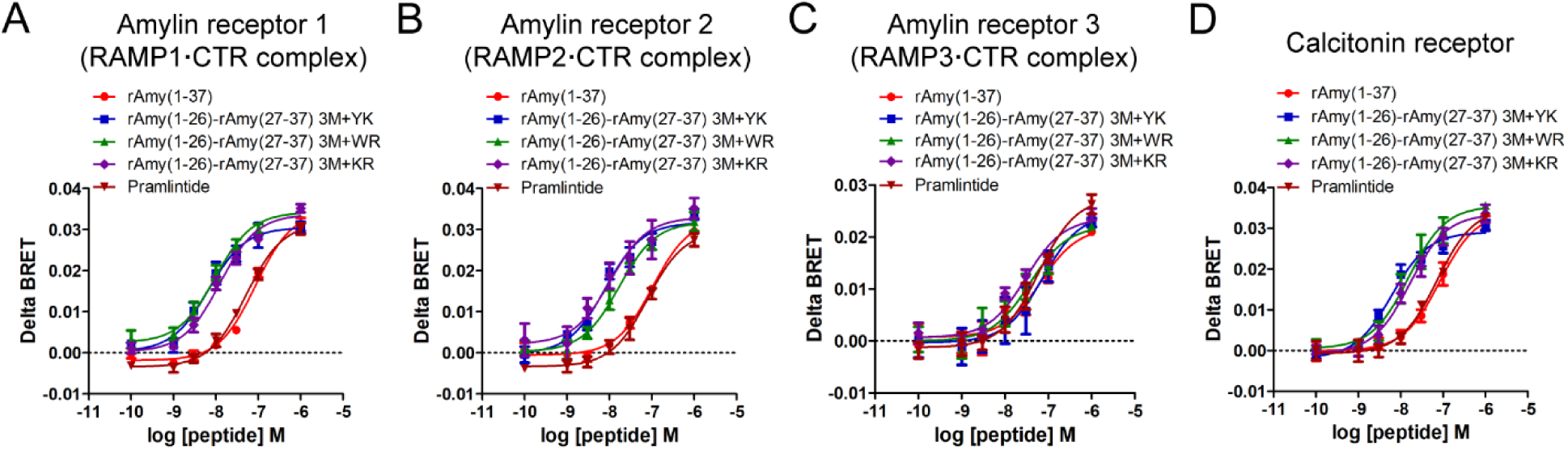
Amylin receptor and calcitonin receptor activation by rat amylin(1–37) analogs with affinity-enhancing mutations for amylin receptor ECD binding. A, B, and C) Amylin receptor 1/2/3 activation by rat amylin(1–37) analogs was measured with G protein recruitment BRET assay. D) Calcitonin receptor activation by rat amylin(1–37) analogs was measured with G protein recruitment BRET assay. Rat amylin(1–37) analogs were generated by introducing 3M + YK, 3M +WR, or 3M + KR mutations. 3M indicates three mutations of rat amylin(27–37): L27W, S34P, and Y37HYP (hydroxyproline). The following two mutations after 3M indicate the mutations at P28 and P29 of rat amylin sequentially. Representative amylin receptor activation curves were shown from at least three independent experiments. rAmy, rat amylin.

## 4. Discussion

### 4.1. Discrepancy between amylin receptor binding affinity and receptor activation

This study found multiple mutations that enhanced peptide ligand affinity for amylin and calcitonin receptor ECDs by over 100-fold (Figure 7, 8, and Table 2). When the affinity-enhancing mutations were incorporated into a full-length thirty-seven amino acid rat amylin peptide, the potency of rat amylin for calcitonin and amylin receptor 1/2 activation was significantly enhanced by up to 10-fold (Figure 9 and Table 3). Affinity enhancement for receptor ECDs appeared to contribute to potency increase for calcitonin and amylin receptor 1/2 activation. However, the degree of enhancement for receptor activation was largely reduced compared to the affinity enhancement for receptor ECDs.

The similar trend for reduced potency enhancement was also observed with calcitonin gene-related peptide (CGRP) and adrenomedullin (AM) receptors. These receptors belong to the class B1 GPCRs where calcitonin and amylin receptors belong to. Booe et al. reported that CGRP N31D/S34P/K35W/A36S mutations increased the CGRP(27–37) fragment affinity for CGRP receptor ECD by more than 1000-fold compared to the affinity of wild-type CGRP(8–37) [28]. Nevertheless, the potency for CGRP receptor activation was enhanced by 4.7-fold with CGRP N31D/S34P/K35W/A36S mutations [28,29]. In addition, AM S48G/Q50W mutations that enhanced peptide affinity for AM receptor ECD by 960-fold showed a 3-fold potency increase for AM receptor activation [29].

These results appear to be involved in the binding mode of peptide hormones for their receptors, in particular class B1 GPCRs. Apparently, the binding affinity for the receptor ECD that is a part of the whole receptor does not represent the activation of the full-length receptor. A recent report with amylin receptor structures clearly showed that the N-terminal part of amylin interacted with the TM domain of the amylin receptor and that the C-terminal part of amylin interacts with amylin receptor ECD [15]. The amylin interaction with amylin receptor ECD is suggested to occur prior to the peptide interaction with TM domain by ‘two domain hypothesis’ [16,30]. Parthier et al. suggested that peptide hormone binding to its receptor ECD triggers the helical formation of the peptide N-terminal part that is crucial for receptor activation [30]. After a peptide hormone binds its receptor ECD, additional steps for receptor TM domain movement appear necessary for the receptor to interact with G protein. It seems apparent that binding to receptor ECD does not represent the whole receptor activation process. This could be the potential reason why discrepancies between the affinity enhancement for receptor ECDs and potency enhancement for the whole receptor activation were observed.

### 4.2. Amylin receptor selectivity among amylin receptor subtypes

The amylin receptor activators designed from this study showed higher receptor activation potency for calcitonin and amylin receptor 1/2, not for amylin receptor 3. Any specific subtype of the amylin receptor has not been reported to play a main role in blood glucose and body weight control. In addition, a clinical drug pramlintide activated all three types of the amylin receptor and the calcitonin receptor with a similar potency when assessed the BRET assay (Figure 9 and Table 3). Cagrilintide that has been tested in clinical trials (AM833) also showed pEC_50_ from 9.71 to 10.14 for cAMP accumulation similarly at calcitonin receptor and amylin 1/3 receptors [31]. Accordingly, it appears that having no selectivity for amylin receptor 1/2 over the calcitonin receptor and activating both amylin and calcitonin receptors with similar potency may not be an issue.

Much less potency change for amylin receptor 3 activation was observed in contrast to the significant 5- to 10-fold potency enhancement shown for amylin receptor 1/2 and calcitonin receptor activation (Figure 9C and Table 3). Although three RAMP types in humans form a complex with CTR TM and ECD domains in the structurally similar way [15], the sequence difference among RAMP types and additional N-glycosylation in RAMP3 may contribute to unique outcome in downstream signaling mediated by amylin receptor 3 activation. RAMP3 has at least three more predicted N-glycosylation sites than RAMP1/2 based on N-glycosylation sequons. In addition, the relatively flattened response in amylin receptor 3-mediated cAMP accumulation compared to CTR alone and amylin receptor 1 was also shown in the previous study [31]. Inconsistency in cell signaling among amylin receptor types was also shown in other studies implicating delicate signaling modulation by RAMPs in amylin receptor activation [32–34].

### 4.3. Targeting amylin N-terminal and middle regions to enhance amylin receptor potency and selectivity

The N-terminal part of amylin interaction with receptor TM domain is known critical for receptor activation. Bower et al. reported several mutations of the N-terminal amylin that significantly increased potency for CTR and/or amylin receptor activation [32]. When human amylin alanine at position 5 was mutated to serine (human calcitonin has serine as the corresponding residue to human amylin alanine at position 5), the mutation increased the potency of human amylin by 14-fold for CTR activation and by 9.1-fold for amylin receptor 3 activation. Q10A, N14A, or V17A mutation of the N-terminal part of human amylin also significantly increased potency for CTR and/or amylin receptor 1 activation. These results indicate that mutagenesis introduced to N-terminal amylin can enhance the potency for calcitonin and amylin receptor activation [32]. The combinational effects of N-terminal mutations and C-terminal affinity-enhancing mutations shown in this study remain to be tested for the future study.

### 4.4. Comparison with cagrilintide and lipidation for future studies

Cagrilintide is a long half-life lipidated amylin receptor activator designed for once-weekly injection [10,31]. Cagrilintide showed 2- to 5-fold potency increases for amylin receptor 1/3 activation (measured as cAMP production) compared to rat amylin potency, but these increases were not statistically significant [31]. The amylin receptor activators designed from this study showed significant 5- to 10-fold potency increases for amylin receptor 1/2 activation compared to rat amylin and pramlintide. Amylin receptor activators designed from this study need to be lipidated to increase pharmacokinetic profile for future studies similarly to the case of cagrilintide. This lipidation may decrease their affinity and potency for amylin receptors. Nevertheless, providing multiple peptide candidates will be useful for obtaining one showing the least change in affinity and potency.

### 4.5. Amylin receptor selectivity over the CGRP receptor

The amylin receptor is the complex of CTR and an accessary protein called RAMP. The calcitonin receptor-like receptor (CLR) also forms the complex with RAMPs [35]. The CLR complex with RAMP1, RAMP2, and RAMP3 was reported as the CGRP receptor, the AM1 receptor, and the AM2 receptor, respectively [36]. The CGRP receptor is a drug target for migraine treatment, and CGRP receptor antagonists and antibodies that block CGRP activity in the human brain are clinically available [37]. CTR and CLR share about 60% amino acid sequence and interestingly, endogenous CGRP could also activate amylin receptor 1 at cell systems [38]. As expected, a CGRP receptor antibody showed antagonism for amylin receptor 1 [39]. Despite significant preference to CGRP receptor antagonism, CGRP receptor antagonists and antibodies could block amylin receptor 1 activation mediated by amylin. Interestingly, a recent clinical report showed that amylin analog pramlintide infusion induced migraine-like attacks in patients similarly to CGRP infusion [40]. This study suggested that amylin receptor agonism may be involved in migraine pathogenesis. Nevertheless, once-a-week injection of an amylin receptor agonist cagrilintide produced headache for 5% to 13% participants of treatment groups that was similar to 12% participants in a placebo-control group [11]. Whether amylin receptor activators generate migraine-like attacks significantly over a placebo-control group should be monitored for future clinical trials.

### 4.6. Pharmaceutical effort on developing amylin receptor activators with a long half-life

Peptide drugs are generally modified to increase bioavailability and to prolong a plasma half-life [41]. Lipidation is one common way to increase the half-life of a peptide drug [42,43]. Plasma albumin interaction with peptide drugs is known to be increased by lipidation. Proper lipidation on peptides contributes to extending a half-life of peptide drugs and successfully enabled once a week injection of peptide drugs with clinical benefits [11,12]. The amylin receptor activators presented in the current study would require lipidation for the increased half-life in animal studies. Lipidation introduced to cagrilintide [10] could be applied to these amylin receptor activators and the modified peptides remain to be tested for amylin receptor activation in a cell system and for anti-obesity efficacy *in vivo*.

### 4.7. Closing remarks

The current study found multiple peptide drug candidates with enhanced activation potency for calcitonin and amylin receptors. Peptide modification and optimization will be pursued prior to testing anti-obesity efficacy in animal studies. Having multiple amylin receptor activators designed by this study can be used as potential candidates for the development of novel anti-obesity peptide drugs.

## Ethics approval and consent to participate

Not applicable

## Consent for publication

Not applicable

## Availability of data and material

Data and materials are available from the corresponding author on reasonable request.

## Author Contributions

SL conceived, designed, and performed experiments and analyzed data. SL wrote the manuscript.

## Acknowledgments

I thank Dr. Nevin Lambert (Medical College of Georgia) for sharing DNA plasmids for BRET assay. I also thank Dr. Scott E. Hemby (High Point University, Fred Wilson School of Pharmacy) for sharing his research equipment I used for this study.

## Abbreviations

sCT: salmon calcitonin
rAmy: rat amylin
CTR: calcitonin receptor
CGRP: calcitonin gene-related peptide AM adrenomedullin
TM: transmembrane domain
ECD: extracellular domain
ECL: extracellular loop

## References

1. Kahn SE, Hull RL & Utzschneider KM (2006) Mechanisms linking obesity to insulin resistance and type 2 diabetes. Nature 444, 840–846.

2. Powell-Wiley TM, Poirier P, Burke LE, Despres J, Gordon-Larsen P, Lavie CJ, Lear SA, Ndumele CE, Neeland IJ, Sanders P, St-Onge M, American Heart Association Council on Lifestyle and Cardiometabolic Health, Council on Cardiovascular and Stroke Nursing, Council on Clinical Cardiology, Council on Epidemiology and Prevention & and Stroke Council (2021) Obesity and Cardiovascular Disease: A Scientific Statement From the American Heart Association. Circulation 143, e984–e1010.

3. Meier JJ (2012) GLP-1 receptor agonists for individualized treatment of type 2 diabetes mellitus. Nat Rev Endocrinol 8, 728–742.

4. Wilding JPH, Batterham RL, Calanna S, Davies M, Van Gaal LF, Lingvay I, McGowan BM, Rosenstock J, Tran MTD, Wadden TA, Wharton S, Yokote K, Zeuthen N, Kushner RF & STEP 1 Study Group (2021) Once-Weekly Semaglutide in Adults with Overweight or Obesity. N Engl J Med 384, 989–1002.

5. Pi-Sunyer X, Astrup A, Fujioka K, Greenway F, Halpern A, Krempf M, Lau DCW, le Roux CW, Violante Ortiz R, Jensen CB, Wilding JPH & SCALE Obesity and Prediabetes NN8022-1839 Study Group (2015) A Randomized, Controlled Trial of 3.0 mg of Liraglutide in Weight Management. N Engl J Med 373, 11–22.

6. Singh G, Krauthamer M & Bjalme-Evans M (2022) Wegovy (semaglutide): a new weight loss drug for chronic weight management. J Investig Med 70, 5–13.

7. Jastreboff AM, Aronne LJ, Ahmad NN, Wharton S, Connery L, Alves B, Kiyosue A, Zhang S, Liu B, Bunck MC, Stefanski A & SURMOUNT-1 Investigators (2022) Tirzepatide Once Weekly for the Treatment of Obesity. N Engl J Med 387, 205–216.

8. Hay DL, Chen S, Lutz TA, Parkes DG & Roth JD (2015) Amylin: Pharmacology, Physiology, and Clinical Potential. Pharmacol Rev 67, 564–600.

9. McQueen J (2005) Pramlintide acetate. Am J Health Syst Pharm 62, 2363–2372.

10. Kruse T, Hansen JL, Dahl K, Schaffer L, Sensfuss U, Poulsen C, Schlein M, Hansen AMK, Jeppesen CB, Dornonville de la Cour C, Clausen TR, Johansson E, Fulle S, Skyggebjerg RB & Raun K (2021) Development of Cagrilintide, a Long-Acting Amylin Analogue. J Med Chem.

11. Lau DCW, Erichsen L, Francisco AM, Satylganova A, le Roux CW, McGowan B, Pedersen SD, Pietilainen KH, Rubino D & Batterham RL (2021) Once-weekly cagrilintide for weight management in people with overweight and obesity: a multicentre, randomised, double-blind, placebo-controlled and active-controlled, dose-finding phase 2 trial. Lancet 398, 2160–2172.

12. Enebo LB, Berthelsen KK, Kankam M, Lund MT, Rubino DM, Satylganova A & Lau DCW (2021) Safety, tolerability, pharmacokinetics, and pharmacodynamics of concomitant administration of multiple doses of cagrilintide with semaglutide 2.4 mg for weight management: a randomised, controlled, phase 1b trial. Lancet 397, 1736–1748.

13. Tilakaratne N, Christopoulos G, Zumpe ET, Foord SM & Sexton PM (2000) Amylin receptor phenotypes derived from human calcitonin receptor/RAMP coexpression exhibit pharmacological differences dependent on receptor isoform and host cell environment. J Pharmacol Exp Ther 294, 61–72.

14. Christopoulos G, Perry KJ, Morfis M, Tilakaratne N, Gao Y, Fraser NJ, Main MJ, Foord SM & Sexton PM (1999) Multiple amylin receptors arise from receptor activity-modifying protein interaction with the calcitonin receptor gene product. Mol Pharmacol 56, 235–242.

15. Cao J, Belousoff MJ, Liang Y, Johnson RM, Josephs TM, Fletcher MM, Christopoulos A, Hay DL, Danev R, Wootten D & Sexton PM (2022) A structural basis for amylin receptor phenotype. Science 375, eabm9609.

16. Hoare SR (2005) Mechanisms of peptide and nonpeptide ligand binding to Class B G-protein-coupled receptors. Drug Discov Today 10, 417–427.

17. Roder C, Kupreichyk T, Gremer L, Schafer LU, Pothula KR, Ravelli RBG, Willbold D, Hoyer W & Schroder GF (2020) Cryo-EM structure of islet amyloid polypeptide fibrils reveals similarities with amyloid-beta fibrils. Nat Struct Mol Biol 27, 660–667.

18. Wan Q, Okashah N, Inoue A, Nehme R, Carpenter B, Tate CG & Lambert NA (2018) PMC5949987; Mini G protein probes for active G protein-coupled receptors (GPCRs) in live cells. J Biol Chem 293, 7466–7473.

19. Aricescu AR, Lu W & Jones EY (2006) A time- and cost-efficient system for high-level protein production in mammalian cells. Acta Crystallogr D Biol Crystallogr 62, 1243–1250.

20. Lee S & Pioszak AA (2020) PMC7686023; Molecular interaction of an antagonistic amylin analog with the extracellular domain of receptor activity-modifying protein 2 assessed by fluorescence polarization. Biophys Chem 267, 106477.

21. Lee S (2021) Development of High Affinity Calcitonin Analog Fragments Targeting Extracellular Domains of Calcitonin Family Receptors. Biomolecules 11, 1364.

22. England CG, Ehlerding EB & Cai W (2016) NanoLuc: A Small Luciferase Is Brightening Up the Field of Bioluminescence. Bioconjug Chem 27, 1175–1187.

23. Lee SM, Hay DL & Pioszak AA (2016) PMC4861438; Calcitonin and Amylin Receptor Peptide Interaction Mechanisms: INSIGHTS INTO PEPTIDE-BINDING MODES AND ALLOSTERIC MODULATION OF THE CALCITONIN RECEPTOR BY RECEPTOR ACTIVITY-MODIFYING PROTEINS. J Biol Chem 291, 8686–8700.

24. Lee SM, Booe JM, Gingell JJ, Sjoelund V, Hay DL & Pioszak AA (2017) PMC5551398; N-Glycosylation of Asparagine 130 in the Extracellular Domain of the Human Calcitonin Receptor Significantly Increases Peptide Hormone Affinity. Biochemistry 56, 3380–3393.

25. Roehrl MH, Wang JY & Wagner G (2004) A general framework for development and data analysis of competitive high-throughput screens for small-molecule inhibitors of protein-protein interactions by fluorescence polarization. Biochemistry 43, 16056–16066.

26. Warnecke A, Sandalova T, Achour A & Harris RA (2014) PyTMs: a useful PyMOL plugin for modeling common post-translational modifications. BMC Bioinformatics 15, 370–6.

27. Lee SM, Jeong Y, Simms J, Warner ML, Poyner DR, Chung KY & Pioszak AA (2020) PMC7225057; Calcitonin Receptor N-Glycosylation Enhances Peptide Hormone Affinity by Controlling Receptor Dynamics. J Mol Biol 432, 1996–2014.

28. Booe JM, Warner ML, Roehrkasse AM, Hay DL & Pioszak AA (2018) PMC5832325; Probing the Mechanism of Receptor Activity-Modifying Protein Modulation of GPCR Ligand Selectivity through Rational Design of Potent Adrenomedullin and Calcitonin Gene-Related Peptide Antagonists. Mol Pharmacol 93, 355–367.

29. Booe JM, Warner ML & Pioszak AA (2020) Picomolar Affinity Antagonist and Sustained Signaling Agonist Peptide Ligands for the Adrenomedullin and Calcitonin Gene-Related Peptide Receptors. ACS Pharmacol Transl Sci 3, 759–772.

30. Parthier C, Reedtz-Runge S, Rudolph R & Stubbs MT (2009) Passing the baton in class B GPCRs: peptide hormone activation via helix induction? Trends Biochem Sci 34, 303–310.

31. Fletcher MM, Keov P, Truong TT, Mennen G, Hick CA, Zhao P, Furness SGB, Kruse T, Clausen TR, Wootten D & Sexton PM (2021) AM833 is a novel agonist of calcitonin family G protein-coupled receptors: pharmacological comparison to six selective and non-selective agonists. J Pharmacol Exp Ther.

32. Bower RL, Yule L, Rees TA, Deganutti G, Hendrikse ER, Harris PWR, Kowalczyk R, Ridgway Z, Wong AG, Swierkula K, Raleigh DP, Pioszak AA, Brimble MA, Reynolds CA, Walker CS & Hay DL (2018) PMC7088893; Molecular Signature for Receptor Engagement in the Metabolic Peptide Hormone Amylin. ACS Pharmacol Transl Sci 1, 32–49.

33. Bailey RJ, Walker CS, Ferner AH, Loomes KM, Prijic G, Halim A, Whiting L, Phillips ARJ & Hay DL (2012) Pharmacological characterization of rat amylin receptors: implications for the identification of amylin receptor subtypes. Br J Pharmacol 166, 151–167.

34. Garelja ML, Bower RL, Brimble MA, Chand S, Harris PWR, Jamaluddin MA, Petersen J, Siow A, Walker CS & Hay DL (2022) Pharmacological characterisation of mouse calcitonin and calcitonin receptor-like receptors reveals differences compared with human receptors. Br J Pharmacol 179, 416–434.

35. McLatchie LM, Fraser NJ, Main MJ, Wise A, Brown J, Thompson N, Solari R, Lee MG & Foord SM (1998) RAMPs regulate the transport and ligand specificity of the calcitonin-receptor-like receptor. Nature 393, 333–339.

36. Hay DL, Garelja ML, Poyner DR & Walker CS (2018) Update on the pharmacology of calcitonin/CGRP family of peptides: IUPHAR Review 25. Br J Pharmacol 175, 3–17.

37. Ogunlaja OI & Goadsby PJ (2022) Headache: Treatment update. eNeurologicalSci 29, 100420.

38. Garelja ML, Walker CS & Hay DL (2022) CGRP receptor antagonists for migraine. Are they also AMY(1) receptor antagonists? Br J Pharmacol 179, 454–459.

39. Garelja ML, Alexander TI, Bennie A, Nimick M, Petersen J, Walker CS & Hay DL (2023) Pharmacological characterisation of erenumab, Aimovig, at two calcitonin gene-related peptide responsive receptors. Br J Pharmacol, 11–21.

40. Ghanizada H, Al-Karagholi MA, Walker CS, Arngrim N, Rees T, Petersen J, Siow A, Morch-Rasmussen M, Tan S, O’Carroll SJ, Harris P, Skovgaard LT, Jorgensen NR, Brimble M, Waite JS, Rea BJ, Sowers LP, Russo AF, Hay DL & Ashina M (2021) Amylin Analog Pramlintide Induces Migraine-like Attacks in Patients. Ann Neurol 89, 1157–1171.

41. Lee MF & Poh CL (2023) Strategies to improve the physicochemical properties of peptide-based drugs. Pharm Res 40, 617–632.

42. Knudsen LB & Lau J (2019) The Discovery and Development of Liraglutide and Semaglutide. Front Endocrinol (Lausanne) 10, 155.

43. Jamaluddin A, Chuang C, Williams ET, Siow A, Yang SH, Harris PWR, Petersen JSSM, Bower RL, Chand S, Brimble MA, Walker CS, Hay DL & Loomes KM (2022) Lipidated Calcitonin Gene-Related Peptide (CGRP) Peptide Antagonists Retain CGRP Receptor Activity and Attenuate CGRP Action In Vivo. Front Pharmacol 13, 832589.

